# The brain-meningeal interface functions as a reservoir and entry site for brain parenchymal macrophages

**DOI:** 10.64898/2026.07.15.738664

**Authors:** Deniz Kaymak, Philip T. Williams, Sabrina Hogan, Marie-Françoise Ritz, Giulia Di Bartolomei, Sihan Zhu, Siling Du, Hassan Melhem, Ana C. Delgado, Tala Shekarian, Marta McDaid, Alexandra Gerber, Igor Smirnov, Patrice Zeis, Timo Schenker, Fiona Gerster, Valerio Sabatino, Zora Baumann, Alexandar Tzankov, Raphael Guzman, Luigi Mariani, Fiona Doetsch, Jonathan Kipnis, Rainer Glass, Gregor Hutter

**Affiliations:** Brain Tumor Immunotherapy and Biology Lab, Department of Biomedicine, University of Basel, University Hospital Basel, Switzerland; Department of Neurosurgery, University Hospital Basel, Switzerland; Department of Neurosurgery, Neurosurgical Research, Ludwig-Maximilians-Universität Munich, Munich, Germany; Brain Immunology and Glia (BIG) Center, Department of Pathology and Immunology, Washington University, St. Louis, MO, USA; Gastroenterology Lab, Department of Biomedicine, University of Basel, University Hospital Basel, Switzerland; Biocenter, University of Basel, Switzerland; Tumor Heterogeneity and Resistance Lab, Department of Biomedicine, University of Basel, Switzerland; Institute of Medical Genetics and Pathology, University Hospital Basel, Switzerland

**Keywords:** Microglia replacement, CNS macrophages, border-associated macrophages, monocyte-derived macrophages, macrophage repopulation, CSF1R, brain-engrafting monocytes, microglial niche, CNS borders, macrophage ontogeny

## Abstract

Microglia are regarded as self-maintaining brain parenchymal macrophages without contribution from adult hematopoiesis. Nevertheless, peripheral macrophage engraftment into the brain has been reported, but the biological variables governing central nervous system (CNS) macrophage niche access remain unclear. We show that CNS macrophage engraftment is determined by the balance between tissue-resident macrophage (TRM) self-renewal and the temporal alignment of niche opening with the availability of engraftment-competent cells and proximity to the vacated niche, rather than prolonged niche vacancy. Fate mapping revealed that monocytes entering the subdural space undergo border-associated macrophage (BAM)-like differentiation, clonal expansion, and transpial migration into the parenchyma, whereas mature BAMs directly repopulate selectively vacated parenchymal niches. We further identify a parenchymal macrophage population with a peripheral BAM-like transcriptional program in aged and neurodegenerative human brains, challenging the dogma of MG-exclusivity. Together, these findings establish a predictive framework for interpreting macrophage maintenance and replacement and for developing macrophage-based therapies.

## INTRODUCTION

The CNS contains anatomically distinct macrophage niches populated by TRMs^1–4^. Among these, microglia (MG) represent the sole macrophage population within the brain parenchyma, whereas BAMs reside at CNS interfaces, including the leptomeninges (LM), perivascular (PV) spaces (subsummarized as subdural BAMs; sdBAMs), the dura mater (duBAMs), and the choroid plexus (CP; cpBAMs). One of the tenets of CNS immune privilege is the restricted accessibility of the brain parenchyma to peripheral immune cells. Hence, MG and sdBAMs are maintained predominantly through local self-renewal^5,6^ without contribution from circulating monocytes under homeostasis^7^. While all CNS macrophage compartments are initially seeded by yolk sac–derived progenitors^2,8–10^, only MG and sdBAMs maintain a predominantly embryonic ontogeny throughout adulthood, whereas cpBAMs and duBAMs undergo partial replacement by bone marrow (BM) derived monocytes, resulting in a mixed ontogeny^2,9,11–13^.

Macrophage depletion and repopulation paradigms have become central tools to interrogate macrophage function and maintenance. However, distinct pharmacological and genetic depletion strategies produce divergent repopulation outcomes, ranging from complete restoration through residual MG self-renewal to extensive peripheral macrophage engraftment^1,7,14–18^. A unifying framework defining the biological principles that govern peripheral macrophage access and CNS niche population, and defines capable cellular sources is currently lacking. Addressing these questions has become important as macrophage replacement therapy is transitioning from preclinical proof-of-concept studies to clinical application^19^. Recent therapeutic success in patients with CSF1R-associated leukoencephalopathy (ALSP) demonstrated that MG replacement could alter disease progression in humans. However, these approaches rely on irradiation- and chemotherapy-based conditioning to create permissive niches for donor-cell engraftment ^14,20^. As the field reaches this translational inflection point, a mechanistic understanding of how CNS macrophage niches are maintained, accessed, and reconstituted will be critical for the development of safer and more effective replacement strategies.

Here, using complementary pharmacological and genetic macrophage depletion models in mice, we systematically dissected the determinants governing CNS macrophage reconstitution and defined physiological principles controlling peripheral macrophage engraftment into the CNS. We identified the coincidence of niche opening and availability of engraftment-competent cells as principal determinants governing competitive occupation of CNS macrophage niches and identified BAMs as spatially privileged reservoirs capable of directly colonizing the MG niche through transpial migration across CNS border interfaces. Finally, we provide evidence that these principles extend to humans by identifying CD163^+^ brain-engrafting peripheral-like macrophages in the aged and neurodegenerating human brain.

## RESULTS

### Monocyte contribution to BAMs is subset-specific at steady state and broadens after niche clearance

To define CNS macrophage identities using a robust and objective framework, we re-analyzed a publicly available single-cell RNA sequencing (scRNA-seq) dataset^1^ originating from adult C57BL/6 mice comprising macrophages isolated from the dura, LM and PV, CP, and brain parenchyma. Unsupervised clustering identified two monocyte populations and six macrophage clusters, each defined by distinct transcriptional programs (**Fig. 1A**), with classical monocytes (CM) expressing high levels of *Ccr2*, *Ly6c2*, and *S100a8*, and non-classical monocytes (NCM) characterized by *Itgal*, *Ace*, and *Ear2* (**Fig. 1A-B**).

**Figure 1:**
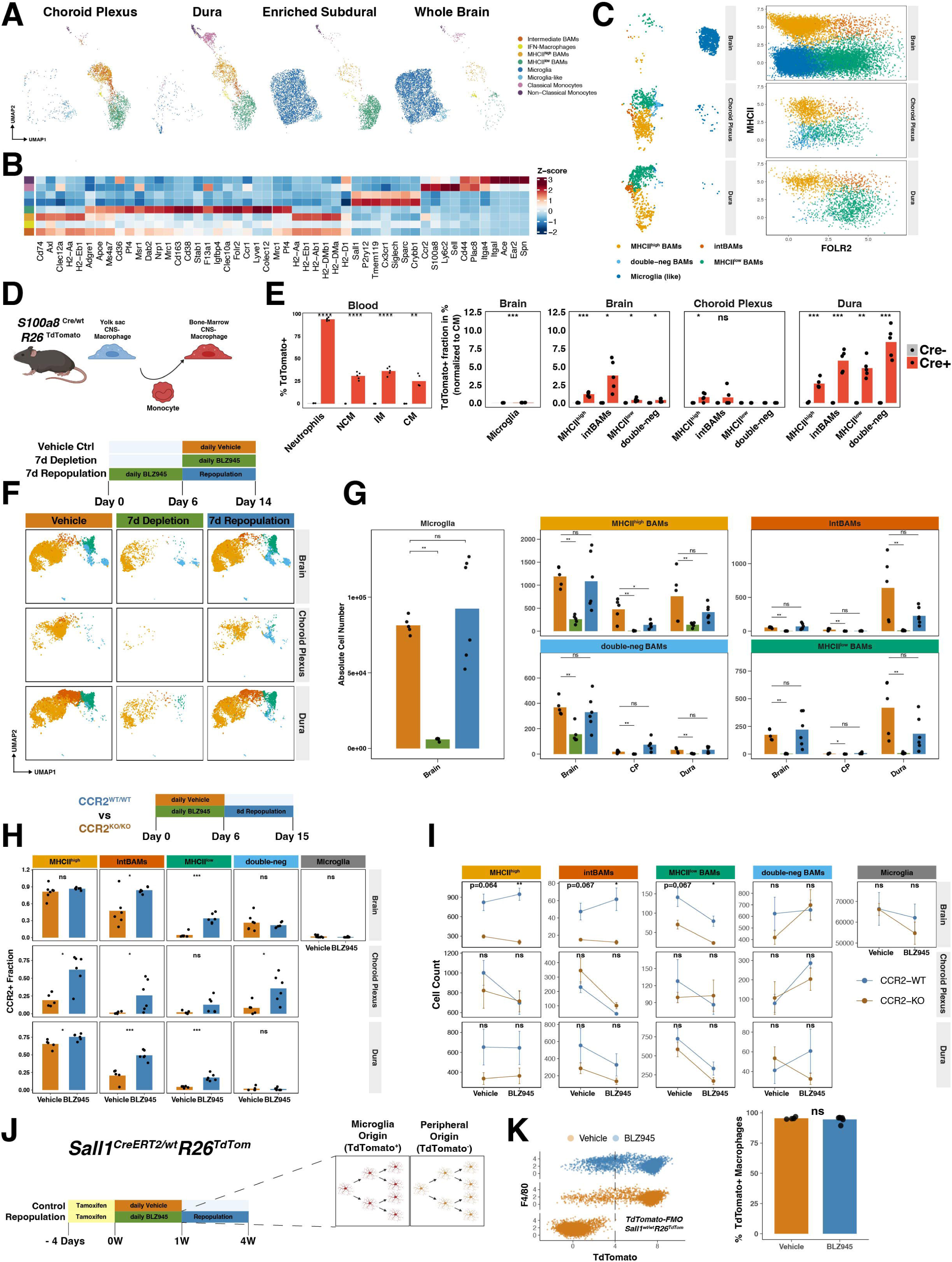
Monocyte contribution to BAMs is subset-specific at steady state and broadens after niche clearance. (**A**) UMAP of re-analyzed scRNAseq dataset^1^ displaying CNS macrophages. (**B**) Heatmap with mean expression of curated BAM−signature marker genes across annotated clusters shown in (A). (**C**) UMAPs and bivariate plots of HDFC analysis of *Sall1^GFP/wt^* mice (n=4 *Sall1^GFP/wt^*, n=5 *Sall1^wt/wt^)*. (**D**) Schematic of *Mrp8^Cre/wt^R26^TdTomato^* mice. (**E**) TdTomato recombination in blood and CNS macrophages. (**F**) Top, experimental outline of depletion/repopulation experiment. Bottom, UMAP of BAMs from brain, CP and dura pre-gated as CD45^+^CD11b^+^Ly6C^-^CD64^+^CD206^+^/Clec12a^+^ macrophages. (**G**) Quantification of clusters in (F) and MG (gated as CD45^+^CD11b^+^Ly6C^-^CD64^+^CD206^-^Clec12a^-^P2y12^+^CD49d^-^). (**H-I**) CNS macrophage repopulation in *Ccr2^wt^* and *CCR2^ko^* mice 8d after BLZ945 treatment. (**H**) Top, experimental scheme. (**H**) Fraction of CCR2-positive macrophages in CCR2^wt^ mice under vehicle and BLZ945 conditions, quantified using the CCR2^ko^-derived thresholds shown in Fig. S1I. See Fig. S1J-K for cluster identity/phenotypes. (**I**) Absolute macrophage counts in *Ccr2^wt^* and *Ccr2^ko^* mice (**J**) Left, experimental schematic of MG lineage-tracing using *Sall1^CreERT2/wt^:R26^TdTomato^* mice. (**K**) Left, representative bivariate plot of brain macrophages. Right, quantification of TdTomato^+^ brain macrophages. Statistical tests: (E, K) Two-sided Welch’s t-tests (G) Pairwise unpaired Wilcoxon rank-sum tests between groups within each tissue and macrophage subset. (H) Linear model per tissue and per BAM subset (CCR2_pos_fraction ∼ treatment) with BH correction. (I) Unpaired two-sided Welch’s t-tests with BH correction.

MG formed a discrete cluster, defined by canonical markers including *P2ry12*, *Tmem119*, *Siglech*, and *Sall1*, confirming their distinct identity relative to all other macrophage populations (**Fig. 1A-B**).

In contrast, BAMs segregated into MHCII^low^ and MHCII^high^ populations, distinguished by reciprocal expression of canonical tissue-resident markers (*Mrc1*, *Lyve1*, *Folr2*, *Pf4*, *Cd163*) and antigen-presentation genes (*H2-Aa*, *H2-Ab1*, *Cd74*) (**Fig. 1A-B**)^1,3,9,21^. We additionally identified an intermediate BAM (intBAMs) population co-expressing MHC class II genes together with markers characteristic of MHCII^low^ BAMs, suggesting a continuum of transcriptional states rather than strictly discrete populations (**Fig. 1A-B, Fig. S1A**).

Consistent with established macrophage life-cycle programs^9^, TLF (including *<u>T</u>imd4*, *<u>L</u>yve1*, *<u>F</u>olr2*)-associated signatures were enriched in MHCII^low^/intBAMs, indicating a long-lived, self-renewing population, whereas CCR2-associated signatures predominated in MHCII^high^ BAMs and monocytes, suggesting ongoing replenishment from CM (**Fig. S1B-C**)

Next, we used cluster-specific signatures to design a high-dimensional flow cytometry (HDFC) panel to identify defined CNS macrophage subsets and guide the selection of subset-specific Cre driver mouse lines for targeted genetic manipulation of these populations *in vivo* (**Fig. 1B**). We confirmed the suitability of these markers for HDFC and histological characterization of CNS macrophage subsets. Specifically, we anatomically dissected cranial dura, CP of the fourth and lateral ventricles as well as brain tissue of MG-specific (*Sall1^GFP^)* reporter mice (**Fig. 1C**). Clustering analysis of macrophages (CD45^+^CD11b^+^Ly6C^-^CD64^+^F480^+^ pre-gated cells) resolved four BAM subsets and one brain-enriched MG population. Projection of cluster identities onto FOLR2/MHCII bivariate plots aligned with the macrophage subsets previously identified by scRNAseq, including MHCII^high^ BAMs, MHCII^high^FOLR2^high^ (= intBAMs), FOLR2^high^MHCII^low^ BAMs (= MHCII^low^ BAMs), and double-negative BAMs. (**Fig. 1C**). IntBAMs were most prominent in the dura mater (**Fig. S1D**). Consistent with scRNAseq, FOLR2^high^ BAMs co-expressed CD38 and CD206 (**Fig. S1E**) so that FOLR2 and CD38 were used interchangeably throughout the study. CCR2 was detectable in MHCII^high^ and intBAMs (**Fig. S1E**). *Sall1*-GFP expression was predominantly confined to the MG/MG-like cluster across tissues, with only low-level expression observed in double-negative BAMs, reflecting gating or clustering ambiguity resulting from the lack of MHCII and FOLR2 expression (**Fig. S1F**). Complementary histological analyses in *Cx3cr1*^GFP/wt^ reporter mice using CD206 and MHCII staining across brain border compartments further supported the anatomical distribution of the identified CNS macrophage populations (**Fig. S1G**).

### Monocytes contribute to BAMs across all CNS borders at steady state and after depletion

ScRNA-seq identified *S100a8* as a highly specific marker for CM. To assess the contribution of hematopoietic stem cell (HSC)-derived monocytes to CNS macrophages, we employed adult *Mrp8*(*S100a8*)^Cre/wt^:*R26*^TdTomato^ mice to HDFC. Cre-mediated recombination labeled neutrophils (∼100%) and ∼25% of circulating CMs, whereas MG remained unlabeled (<0.04%), confirming the model’s specificity (**Fig. 1D-E**). We detected low but measurable TdTomato recombination across BAM populations (0–10% depending on the subset) with the most consistent monocyte contribution to intBAMs in brain and dura (**Fig. 1E**).

Colony-stimulating factor 1 receptor (CSF1R) signaling is required for the survival and maintenance of macrophages^22^, and its pharmacological inhibition induces their rapid depletion^1,23,24^. To examine CSF1R-dependency and the niche recovery of CNS macrophages following pharmacological depletion, we treated adult mice with the CSF1R inhibitor (CSF1Ri) BLZ945 for 7d, followed by immediate HDFC analysis or a 7d recovery period after treatment withdrawal (**Fig. 1F-G, Fig. S1H**). CSF1R inhibition resulted in efficient depletion of all CNS macrophage populations across CNS compartments. Upon treatment discontinuation, MG and BAMs in the brain fully repopulated within 7d comparable to vehicle-treated controls. In contrast, cpBAM and duBAMs trended towards incomplete repopulation, suggesting a lower intrinsic self-renewal capacity than MG (**Fig. 1G**).

To assess the monocyte dependency of CNS macrophage niche replenishment, we compared repopulation following CSF1Ri in *Ccr2^wt^*and *Ccr2^ko^* mice, as CCR2 is required for egress of CM from the BM and their tissue recruitment^25^ (**Fig. 1H-I**). Eight days after CSF1Ri withdrawal, we observed an increased fraction of CCR2-expressing BAMs across all border compartments in CCR2^wt^ mice (**Fig. 1H**, see **Fig. S1I for CCR2-thresholds, Fig. S1J** for UMAP and **Fig. S1K** for CCR2-KO confirmation), suggesting recruitment of CCR2^+^ monocyte-derived macrophages (MDMs) into the recovering niche. In contrast, MG showed no CCR2 expression or differential abundance between genotypes (**Fig. 1H-I**). MHCII^high^ BAMs (including intBAMs) across brain and dura trended towards reduced cell numbers at baseline (vehicle) in CCR2^ko^ mice. However, only BAMs in the brain displayed a significant decrease in repopulation across all (except double-negative cells) subsets (**Fig. 1I**). These findings indicate that BAM repopulation, including MHCII^low^subsets, involved CCR2+ macrophages derived from CM. While cpBAMs and duBAMs repopulated independently of CCR2, access to the subdural compartment depended on CCR2.

To determine whether monocytes enter the niche-depleted brain parenchyma, we combined CSF1Ri with MG-specific fate-mapping *Sall1^CreERT2/wt^:R26^TdTomato^* reporter mice. *Sall1* is an embryonically restricted MG marker not expressed by BAMs nor MDMs^18,26,27^. Animals were pretreated with tamoxifen for 5d to induce TdTomato expression in MG, followed by 7d of BLZ945 administration and a 3-week repopulation period (**Fig. 1J**). Repopulating macrophage analysis revealed uniform retention of TdTomato labeling, demonstrating their origin from pre-existing MG (**Fig. 1K**).

Together, these data established that MG self-renew following pharmacological CSF1R-mediated niche depletion without contribution from CM. In contrast, BAMs across all extra-parenchymal CNS compartments, exhibited constitutive engraftment by MDMs, which was further enhanced after niche depletion. While du- and cpBAMs repopulated in a CCR2-independent manner, sdBAM renewal depended on CCR2. However, the brain parenchyma remained refractory to MDM engraftment, reflecting a combination of restricted niche access and the robust self-renewal of MG.

### CSF1R and PU.1 control differentiation of Ly6C^high^ blood monocytes into downstream subsets while being dispensable for their maintenance

The failure of CM to engraft the MG-depleted brain following CSF1Ri prompted us to investigate whether CSF1R signaling regulates monocyte maintenance or differentiation competence. Given conflicting reports regarding monocyte CSF1R dependency^28–37^, we combined pharmacological and genetic approaches to systematically dissect its role.

Murine monocytes encompass classical (CM; Ly6C^high^CCR2^high^CD43^low^) and non-classical (NCM; Ly6C^-^CD43^high^) subsets^38–40^, with CMs progressively differentiating into NCMs through an intermediate monocyte state (IM; Ly6C^int^CD43^int^)^40,41^, all of which express CSF1R^42^.

We performed a time-resolved analysis of monocyte subsets during CSF1R inhibition and recovery (Fig. 2A). Blood was collected during the depletion (d5 and 7) and recovery phase (d9 and 11). CSF1Ri rapidly depleted IMs and NCMs, whereas CM numbers remained stable (**Fig. 2B-C**). Following CSF1Ri withdrawal, IMs rebounded within 2d and transiently exceeded control levels. NCM recovery lagged behind the IM response and followed the transient IM peak, reaching levels above controls by d11 (**Fig. 2C**). These kinetics are consistent with sequential CM-to-IM-to-NCM reconstitution and rapid recovery of differentiation-competent monocytes after CSF1Ri withdrawal.

**Figure 2:**
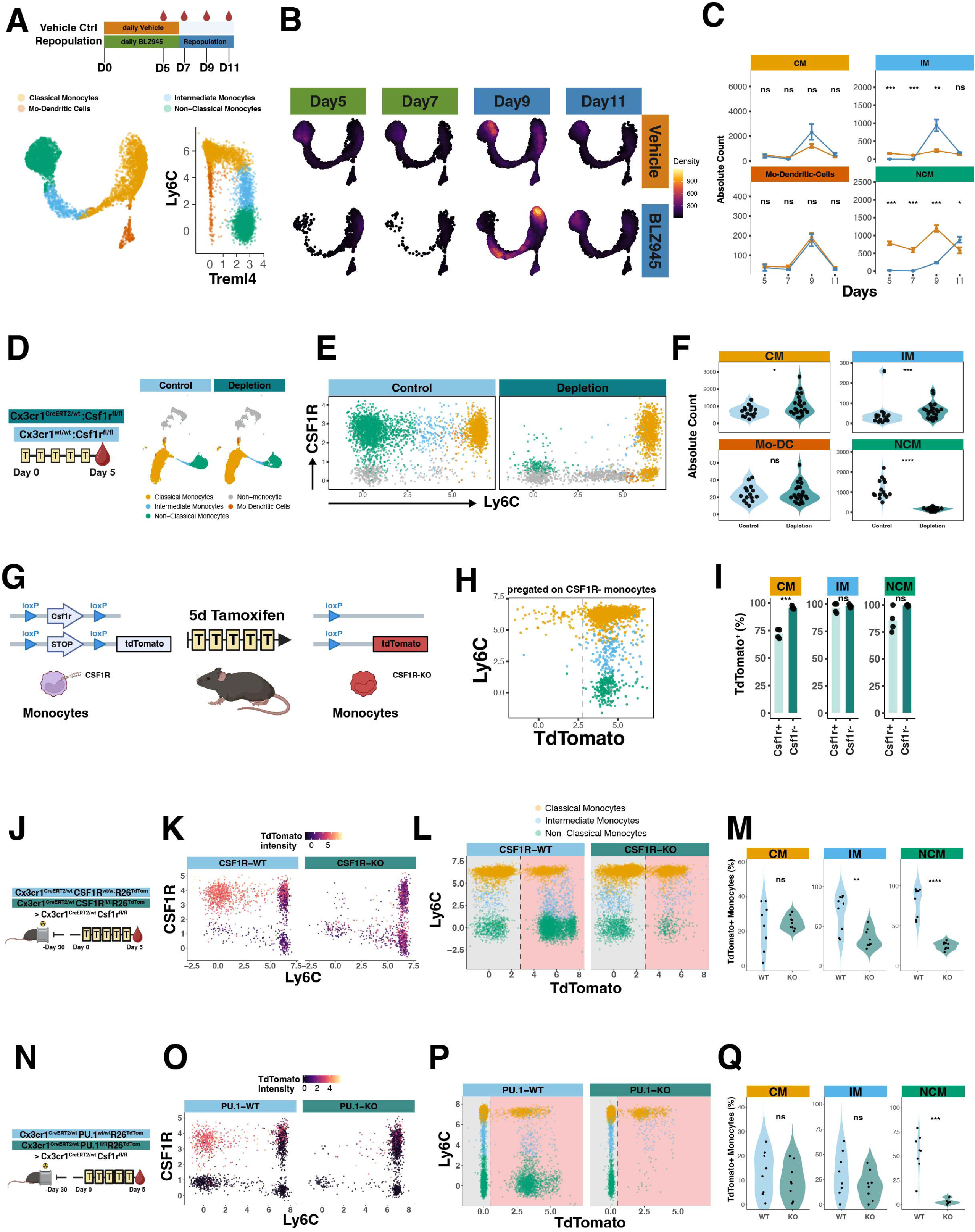
CSF1R and PU.1 control differentiation of Ly6C^high^ monocytes into downstream subsets while being dispensable for their maintenance. (**A**) Top, experimental scheme: *Flt3*^Cre/wt^:*R26*^TdTomato^ mice (n = 4 mice/group). Bottom, UMAP and cluster-identity projected onto bivariate plot (**B**) UMAP-based density plot of monocyte subsets. (**C**) Longitudinal quantification of absolute monocyte counts. (**D**) Left, experimental scheme of *Cx3cr1^CreERT2/wt^:Csf1r^fl/fl^* (n=22) vs *Cx3cr1^wt/wt^:Csf1r^fl/fl^* (n=15). Right, UMAP of CD11b^+^/singlets/live/NK1.1^-^Ly6G^-^Siglec-F^-^ cells. (**E**) Projection of cluster identities in (D) onto bivariate plot. (**F**) Absolute counts of monocytes. (**G**) Schematic of *Cx3cr1*^CreERT2/wt^:*Csf1r*^fl/fl^:*R26*^TdTomato^ mice (n=4). (**H**) TdTomato expression in CSF1R□ monocytes with (**I**) quantification of the TdTomato□ fraction among CSF1R□ and CSF1R□ populations. (**J-M**) Head-shielded BM chimeras with reconstitution of *Cx3cr1*^CreERT2/wt^:*Csf1r*^fl/fl^ recipients with either TdTomato^+^ CSF1R-WT or CSF1R-KO BM. (**J**) Experimental scheme. (**K**) Representative FC plots of blood monocytes. (**L**) FC plots with cluster-based annotation (see Fig. S2N) of monocyte subsets showing gating of TdTomato□ monocytes. (**M**) Fraction of TdTomato□ monocytes per condition (n=9 CSF1R-WT, n=9 CSF1R-KO). (**N–Q**) As in (J–M), but using TdTomato⁺ PU.1-WT or PU.1-KO donor BM (n=8 PU.1-WT, n=8 PU.1-KO). See Fig. S2P for clustering. Statistical tests: (C) Two-sided unpaired t-tests with BH correction (F) Linear mixed-effects model followed by estimated marginal means pairwise comparisons with BH correction. (I) Two-sided paired t-tests. (M, Q) Two-sided unpaired t-tests.

To determine whether apparent preservation of CM reflected reduced CSF1R dependency or continuous BM replenishment, we analyzed monocyte subsets (d5 in **Fig. 1H)** in CSF1Ri-treated Ccr2^⁻/⁻^ mice, in which BM egress of Ly6C^high^ monocytes is impaired^25^. As expected, *Ccr2*^-/-^monocytes lacked CCR2 expression (**Fig. S2A-B,** see UMAP in **Fig. S2A** for cluster identity) resulting in a marked reduction of CM in vehicle treated mice (**Fig. S2C-D**). Notably, BLZ945 treatment did not result in further depletion of CM in *Ccr2*^-/-^ animals. In contrast, IM and NCM were robustly depleted in both genotypes (**Fig. S2E**). Furthermore, among CM-subsets (defined by the expression of CD319 and CD177^43^), no preferential depletion was observed in BLZ945 treated *Ccr2*^-/-^ mice (**Fig. S2F-G**). These findings argue against compensatory replenishment as an explanation for CMs persistence.

To overcome the constraints of off-target kinase inhibition elicited by BLZ945^44^, we next employed a *Cx3cr1*-driven inducible genetic mouse model. Having confirmed *Cx3cr1* expression and efficient recombination across BM and all monocyte subsets in *Cx3cr1*^CreERT2/wt^:*R26*^TdTomato^ and *Cx3cr1*^GFP^ mice (**Fig. S2H-J**), we generated *Cx3cr1*^CreERT2/wt^:*Csf1r*^fl/fl^ animals to enable conditional deletion of *Csf1r*. Following tamoxifen-induced deletion (**Fig. 2D**), blood HDFC analysis revealed a loss of CSF1R-expressing monocyte subsets. NCMs appeared nearly absent, and IMs were strongly reduced. Concurrently, we observed distinct Csf1r^-^ Ly6C^high^ and Ly6C^int^ populations, indicating that loss of CSF1R expression might not coincide with loss of the underlying cells (**Fig. S2K**).

To resolve monocyte identities independently of CSF1R expression, we performed CSF1R-independent unsupervised clustering (**Fig. 2D)** and marker-based annotation (**Fig. S2L**). Csf1r^-^ cells (**Fig. S2M**) clustered together with their respective counterparts in control mice, confirming preservation of monocyte identity (**Fig. 2D**). Projection of cluster identity onto conventional bivariate plots demonstrated that CSF1Rr^-^Ly6C^high^ cells corresponded to bona fide CMs that have lost surface CSF1R. Similar CSF1R^-^ cells were observed across IMs and NCMs clusters (**Fig. 2E**). Differential abundance analysis showed near-complete depletion of NCMs, whereas IMs and CMs were increased (**Fig. 2F**).

To validate the monocyte identity of CSF1R-deficient cells, we used *Cx3cr1*^CreERT2/wt^:*Csf1r^f^*^l/fl^:*R26*^TdTomato^ reporter mice (**Fig. 2G**). Following 5d of tamoxifen treatment, CSF1R^-^ cells uniformly expressed TdTomato across all monocyte subsets (**Fig. 2H-I**) confirming their monocyte identity.

To determine whether loss of CSF1R expression resulted in monocyte cell death or persistence, we next assessed their survival dependency. Specifically, we employed a competitive BM chimera approach to probe monocyte blood niche occupancy and replenishment dynamics. *Cx3cr1*^CreERT2/wt^:*Csf1r^f^*^l/fl^ recipient mice were head-shield irradiated and reconstituted with TdTomato^+^ BM cells (**Fig. 2J**) from either CSF1R-WT (*Cx3cr1^CreERT2/wt^:Csf1r^wt/wt^:R26^TdTomato^*) or CSF1R-KO (*Cx3cr1^CreERT2/wt^:Csf1r^fl/fl^:R26^TdTomato^*) mice. CSF1R-independent survival was inferred from stable donor chimerism, whereas depletion was indicated by preferential replacement with WT donor cells. In this setting, NCMs were efficiently depleted (**Fig. S2N-O**) and failed to recover in CSF1R-KO donor chimeras, whereas they were efficiently replenished by CSF1R-WT donor cells (**Fig. 2K-M**). IMs showed a similar but less pronounced pattern. In contrast, although CMs numbers increased (**Fig. S2P**), they did not show differential donor contribution between CSF1R-WT and -KO conditions, indicating that no replenishable niche was generated in the blood (**Fig. 2K-M**).

To determine whether this subset-specific dependency reflects a broader requirement for myeloid differentiation programs, we performed analogous experiments targeting the transcription factor *Pu.1* (**Fig. 2N**). Consistent with the CSF1R model, PU.1-KO donor cells failed to replenish the NCM population, whereas PU.1-WT donor cells efficiently restored it (**Fig. 2O-Q**) and CMs remained unaffected (**Fig. S2P-Q**).

Collectively, CSF1R defined a differentiation checkpoint within the monocyte trajectory. Loss of CSF1R signaling led to IM and NCM depletion while preserving CMs, indicating a block in subset progression rather than uniform depletion. Despite the rapid restoration of differentiation competence after CSF1Ri, CM failed to outcompete the rapid MG-mediated brain repopulation.

### Impaired self-renewal of TRMs enables monocytes to replace all CNS macrophages, including microglia and subdural BAMs

Building on the observations that both MG competitive dominance and the intrinsic properties of monocytes shape the ability of CMs to differentiate and engraft the CNS, we next examined whether reducing MG competitiveness was sufficient to shift this balance and permitted monocyte access to CNS macrophage niches.

By inducing a permanent genetic ablation of *Csf1r* in *Cx3cr1*^CreERT2/wt^:*Csf1r*^fl/fl^ mice, we reduced the pool of proliferative MG capable of replenishing the parenchymal niche. Following a 7d tamoxifen-mediated depletion phase and a subsequent 25d replenishment period, we observed robust engraftment of CLEC12A⁺F4/80^high^ cells consistent with previously reported peripheral macrophage signatures^18^ (**Fig. S3A**).

To determine their origin and distribution, we generated head-shield chimeras by transplanting TdTomato-labeled BM (*Cx3cr1*^CreERT2/wt^:*R26*^TdTomato^) into *Cx3cr1*^CreERT2/wt^:*Csf1r*^fl/fl^ and *Cx3cr1*^CreERT2/wt^:*Csf1r*^wt/wt^ control mice, enabling selective tracing of MDMs (**Fig. 3A**). Head shielding was employed to avoid irradiation-induced conditioning of the CNS, which can promote inflammatory monocyte recruitment and engraftment^7,45–47^. Dosimetric assessment of the head-shielding device confirmed 95-99% head radioprotection, while allowing body irradiation (**Fig. S3B**). In the absence of niche depletion, donor-derived cells were restricted to CNS border compartments, including sdBAMs (**Fig. 3B-C and Fig. S3C**). In contrast, following genetic niche depletion and repopulation for 25d, TdTomato⁺ MDMs replaced all CNS macrophages in the dura, CP, subdural compartment, and the brain parenchyma (**Fig. 3B-C**).

**Figure 3:**
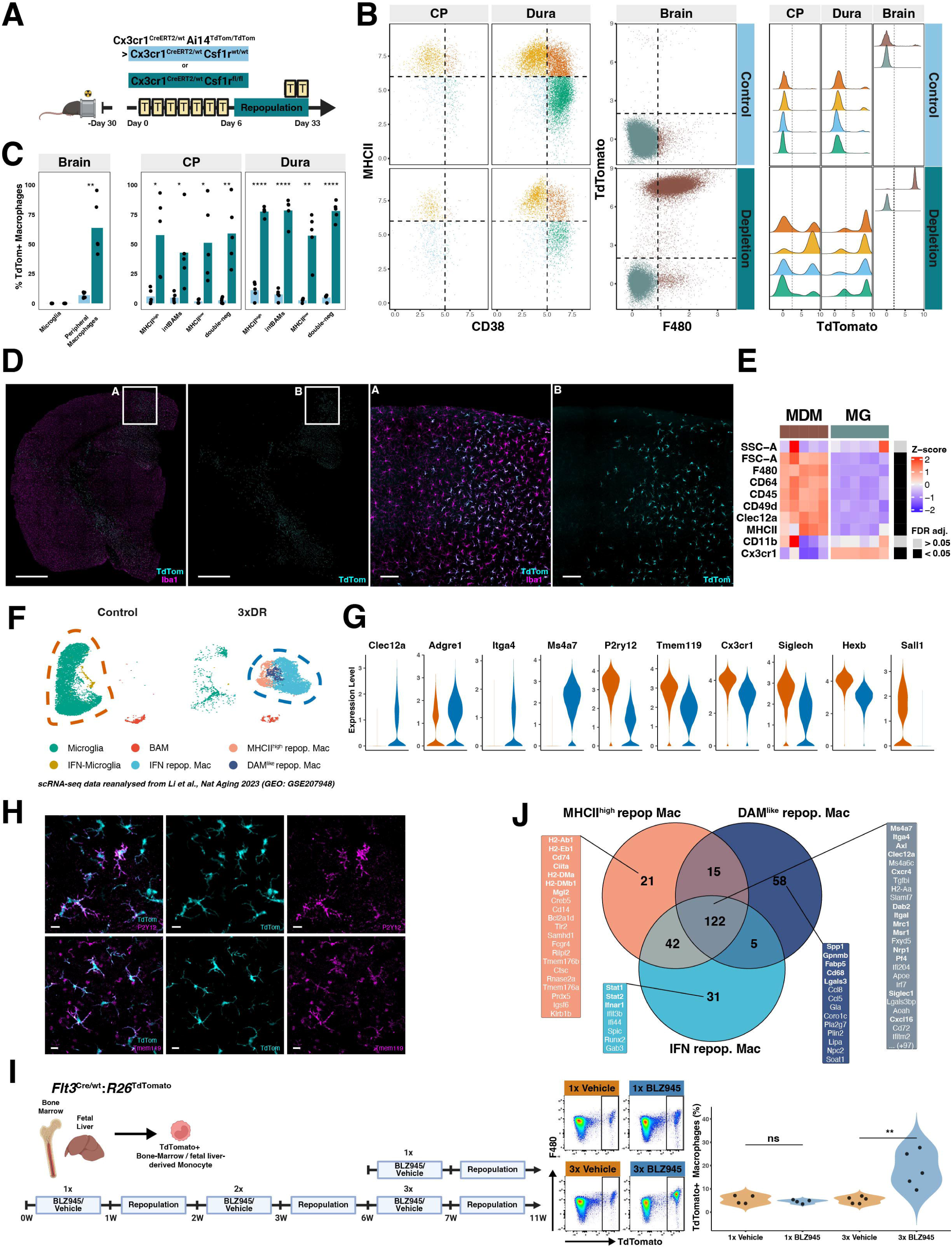
Impaired self-renewal of TRMs enables monocytes to replace all CNS macrophages, including microglia and subdural BAMs. (**A**) Experimental scheme of head-shield chimeras. *Cx3cr1*^CreERT2/wt^:*Csf1r*^fl/fl^ or *Cx3cr1*^CreERT2/wt^:*Csf1r*^wt/wt^ mice were reconstituted with *Cx3cr1*^CreERT2/wt^:*R26*^TdTomato^ BM, followed by tamoxifen-induced recombination and 4-week repopulation. TdTomato recombination was re-induced 2 days before tissue collection. (**B**) Representative gating strategy for CNS macrophages in CP, dura, and brain. Ridge plots show TdTomato expression of gated populations used for quantification (**C**) Percentage of TdTomato^+^ macrophages as gated in (B) (n=5 depletion, n=5 control, one experiment. (**D**) Representative sections of mouse brains from mice in (A-C), stained for Iba1. Scale bars, 1mm, 100 µm. (**E**) Heatmap of Z-score scaled median fluorescence intensity in peripheral macrophages from depletion mice and MG from control mice as defined in (B). (**F**) UMAP of the re-analyzed published 3xDR paradigm by^49^ (n=3 control, n=2 3xDR). See Fig. S3E (**G**) Violin plots highlighting key differentially expressed genes between circled clusters in (F). (**H**) Representative brain sections of head-shield chimera mice from experiment in (A-E) stained for P2Y12 and TMEM119. Scale bar, 10 µm. (**I**) Experimental scheme of repetitive (1xDR, 3xDR and controls) CSF1Ri experiment using Flt3^Cre/wt^R26^TdTomato^ mice. Middle, representative gating of peripheral macrophages (pre-gating in **Fig. S3G**) used for quantification of percentages TdTomato+ macrophages in brains (right). n=4 1xVehicle, n=4 1xDR, n=5 3xVehicle and n=5 for 3xDR, one experiment. (**J**) Venn diagram of shared and uniquely expressed genes among repopulating macrophages. Statistical tests: (C, I) Unpaired two-sided Welch’s t-test. (E) Unpaired two-sided t-tests with BH correction.

Histological analysis confirmed the parenchymal localization of TdTomato^⁺^ repopulating macrophages (**Fig. 3D**), which retained a peripheral macrophage phenotype marked by F4/80, CLEC12A, MHCII, and AXL expression (**Fig. 3E and Fig. S3D**). Together, these data indicated that MDMs contributed continuously to BAM turnover, with increased replacement following depletion. However, access to the parenchymal microglial niche required sustained disruption of TRM occupancy.

### Exhaustion of MG self-renewal enables MDM brain engraftment

Repeated cycles of pharmacological MG depletion have been reported to delay niche refilling^48^, suggesting that the MG proliferative capacity may become limited over time. In a further study^49^, employing a similar iterative depletion paradigm, repopulating macrophages were analyzed by scRNAseq and interpreted as “aged” MG based on the expression of *P2yr12* and *Tmem119*^48,49^. We hypothesized that the delayed repopulation following iterative CSF1R-mediated depletion would create a permissive window for brain engraftment by MDMs.

Reanalysis of the published scRNAseq dataset^49^ (**Fig. 3F-G, Fig. S3E** for experimental outline) revealed elevated expression of *Clec12a*, *Adgre1* (F4/80), *Itga4* (CD49d), and *Ms4a7*, with a reduced expression of core MG genes including *P2ry12* and *Tmem119*, as well as complete absence of *Sall1* expression in repopulating macrophages (**Fig. 3G**). Importantly, comparing these results with data obtained from our head-shield chimera model (**Fig. 3A-E**), we found that TdTomato^+^ brain-engrafting MDMs similarly upregulated canonical MG markers such as P2Y12 and TMEM119 upon brain parenchyma entry (**Fig. 3H**), demonstrating that marker expression alone is insufficient to define cellular origin. This suggests that the repopulating cells observed in iterative depletion models reflect MDMs rather than aged MG. To test this, we combined iterative CSF1Ri (one and three times depletion/repopulation = 1xDR and 3xDR) with monocyte fate mapping using *Flt3*^Cre/wt^:*R26*^TdTomato^ mice, which label monocyte-derived cells with TdTomato (**Fig. 3I**; monocyte recombination shown in **Fig. S3E**). While 1xDR failed to increase brain influx of TdTomato⁺ MDMs compared to the vehicle-treated group, 3xDR led to a marked increase in brain-engrafting MDMs within the fully repopulated brain (**Fig. 3I**, gating shown in **Fig. S3G)**.

Notably, monocyte differentiation was not impaired by repeated depletion, as we observed no decrease in numbers of CM and their downstream subsets across depletion conditions (**Fig. S3H-J**). Thus, while monocyte availability and differentiation capacity were preserved, engraftment was only observed under conditions in which MG self-renewal was progressively impaired.

ScRNAseq analysis^49^ identified multiple MDM transcriptional states, including MHCII^high^ (*H2-Ab1, Cd74, Ciita, Mgl2*), disease-associated macrophage-like (DAM-like; *Spp1, Gpnmb, Fabp5, Cd68, Lgals3*), and interferon-response (IFN repop. Mac; *Stat1, Stat2, Ifnar1*) populations (**Fig. 3F,J**, see **Fig. S3K**). Despite this heterogeneity, all clusters shared a core transcriptional signature (*Ms4a7, Itga4, Axl, Clec12a, Itgal, Mrc1, Fxyd5, Pf4, H2-Aa*) distinct from MG and consistent with MDM origin (**Fig. 3J**).

### The presence of engraftment competent cells during niche opening overrides the requirement for prolonged niche vacancy

While both CSF1Ri and Cx3cr1-driven Csf1r deletion simultaneously perturbed TRMs and the recruitable monocyte pool, we sought to experimentally decouple niche opening from the availability of engraftment-competent cells. We generated head-shield chimeras in which *Cx3cr1*^CreERT2/wt^:*Csf1r*^fl/fl^ recipient mice were reconstituted with either CSF1R-WT (*Cx3cr1*^CreERT2/wt^:*Csf1r*^wt/wt^:*R26*^TdTomato^) or CSF1R-KO (*Cx3cr1*^CreERT2/wt^:*Csf1r*^fl/fl^:*R26*^TdTomato^) TdTomato-labeled BM cells (**Fig. 4A**). At 1d after tamoxifen treatment, TdTomato^⁺^ macrophages in CSF1R-WT chimeras rapidly repopulated the dura, CP, and brain parenchyma, whereas CSF1R-KO monocytes contributed minimally to macrophage repopulation in these compartments (**Fig. 4A–B**). Confocal imaging of the dorsal brain surface demonstrated robust engraftment of MDMs within the LM of CSF1R-WT chimeras but not CSF1R-KO chimeras (**Fig. 4B–C**). Moreover, CSF1R-WT-derived MDMs exhibited morphological features consistent with monocyte-to-macrophage transition, including increased cell area and perimeter together with reduced circularity, compared with the rare CSF1R-KO cells (**Fig. 4D**). Moreover, MDMs in CSF1R-WT conditions displayed an absolute enrichment of both MHCII^high^ and MHCII^low^ MDMs, while CSF1R-KO MDMs appeared arrested in the MHCII^high^ state, with nearly no MHCII^low^ MDMs detected (**Fig. 4E**), consistent with the differentiation block observed in (**Fig. 2J-Q**) and previously described monocyte differentiation trajectories^40,50^.

**Figure 4:**
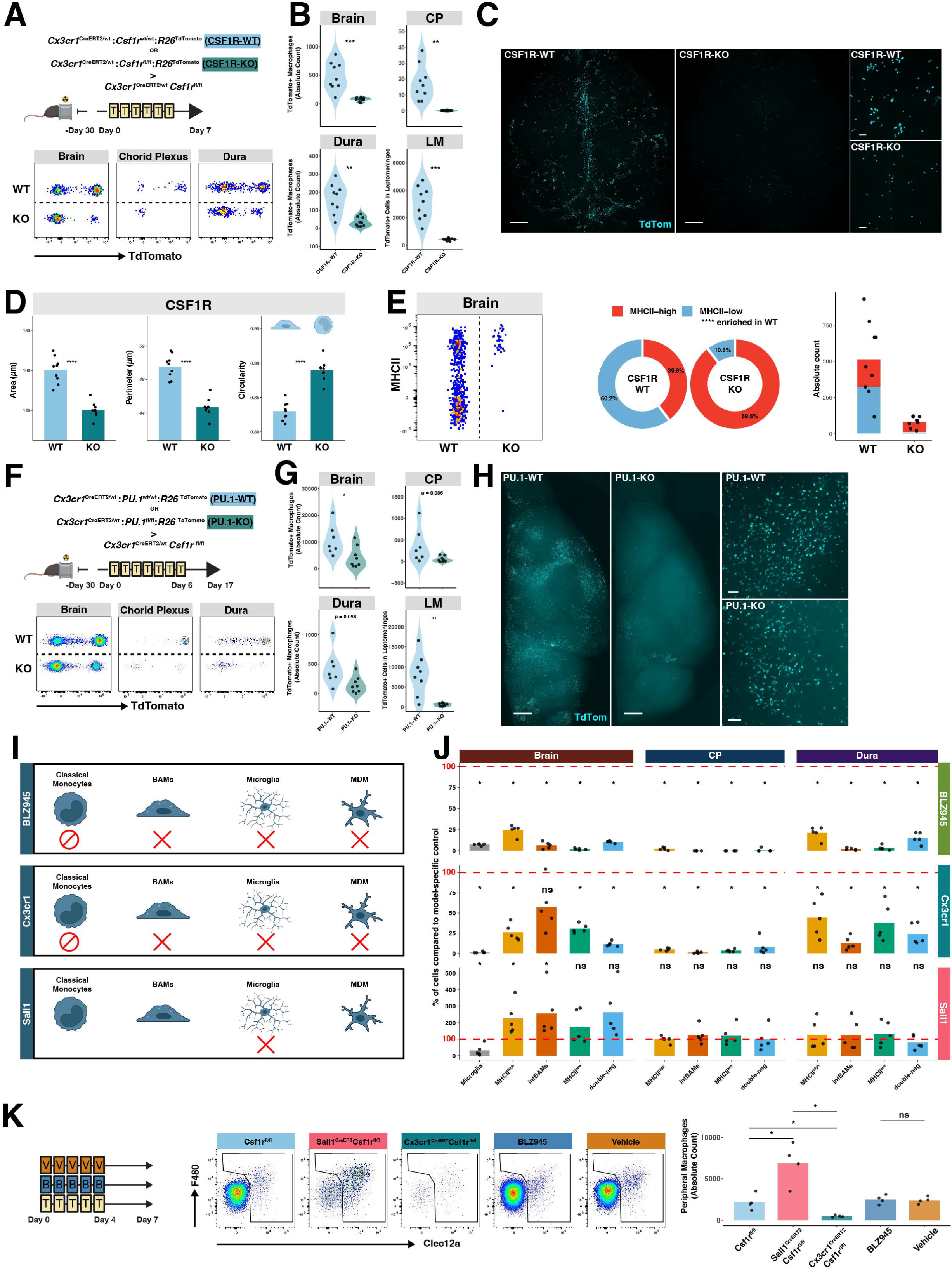
The presence of engraftment competent cells during niche opening overrides the requirement for prolonged niche vacancy. (**A**) Experimental design of head-shielded BM chimeras reconstituted with CSF1R-WT or CSF1R-KO donor BM. Bottom, representative gating of TdTomato⁺ macrophages. (**B**) Absolute count of TdTomato^+^ macrophages by HDFC for brain, CP and dura. Leptomeningeal count was determined by confocal imaging shown in (C). n=9 CSF1R-WT and n=9 CSF1R-KO, one experiment. (**C**) Representative en face confocal imaging of the dorsal brain surface (Scale bar, 1mm, 100 µm) with (**D**) quantification of cell area, perimeter, and circularity across conditions. (**E**) Left, representative FC plot of MHCII expression among TdTomato^+^ repopulating macrophages in brains in (A). Middle, fractional composition of MHCII^+^ and MHCII^-^ cells with absolute counts displayed (right). (**F-H**) As in (A-C), but using TdTomato⁺ PU.1-WT or PU.1-KO donor BM (n=8 PU.1-WT, n=8 PU.1-KO), 7d tamoxifen treatment and tissue harvest on d17 (**I**) Schematic overview on depletion model differences, cross=depleted, stop sign=differentiation block. (**J**) Side-by-side comparison of macrophage depletion across non-selective and microglia-selective depletion models in brain, CP, and dura. Depletion is shown as fraction of the model-specific control mean (= dashed red line), normalized to 100% (vehicle-treated mice for BLZ945; *CreERT2^wt/wt^* tamoxifen-treated mice for depletion models). Analysis after 7d BLZ945 treatment or 5d following tamoxifen-induced CSF1R deletion in genetic models. (**K**) *Left*, experimental scheme comparing early repopulation across models, n=4/group. Mice received tamoxifen (T) from d0-4, alternatively BLZ945 (B) or vehicle (V). Analysis 2d after treatment. *Middle*, Representative gating of CD45^+^CD11b^+^Ly6C^-^Ly6G^-^NK1.1^-^CD64^+^F480^high^Clec12a^high^ peripheral macrophages and (right) absolute counts. Statistical tests: (B, D, E, G) Unpaired two-sided Welch’s t-test. (J) Two-sided unpaired Wilcoxon rank-sum tests with BH correction using absolute counts. (K) Pairwise unpaired two-sided Welch’s t-tests.

Importantly, the number of TdTomato^+^ CMs in the blood was even increased in CSF1R-KO recipient mice (**Fig. S4A**).

To further validate the role of engraftment competence independently of CSF1R signaling, we employed an orthogonal genetic approach targeting the transcription factor PU.1, a master regulator of myeloid differentiation. Head-shielded *Cx3cr1*^CreERT2/wt^:*Csf1r*^fl/fl^ recipient mice were reconstituted with either PU.1-KO (*Cx3cr1*^CreERT2/wt^:*PU.1^fl^*^/fl^:*R26*^TdTomato^) or PU.1-WT (*Cx3cr1*^CreERT2/wt^:*PU.1*^wt/wt^:*R26*^TdTomato^) TdTomato-labeled BM cells. Following tamoxifen-induced depletion for 7d and 10d recovery, CNS compartments were analyzed at 17d (**Fig. 4F**).

Consistent with our previous findings, circulating CMs number were comparable between conditions (**Fig. S4B**). PU.1-WT donor monocytes robustly repopulated macrophage compartments across the dura, LM, and brain parenchyma (see gating strategy and morphological changes in **Fig. S4C-D**), whereas PU.1-KO CMs showed reduced contributions (**Fig. 4F-H**).

In parallel, we sought to employ a MG-specific depletion strategy to preserve CMs. *Sall1* is neither expressed in CM (**Fig. S4E-F**) or BAMs (**Fig. S1F**) nor induced upon CNS entry and differentiation of brain-engrafting MDMs (**Fig. 3G**). Consequently, *Sall1*^CreERT2/wt^:*Csf1r*^fl/fl^ mice enable selective MG depletion (model differences illustrated in **Fig. 4I**). While CSF1Ri treatment and Cx3cr1-driven CSF1R deletion induced widespread depletion of CNS macrophages, *Sall1*-driven depletion did not reduce BAM numbers across CNS compartments but resulted in a significantly increased abundance of MHCII^high^ sdBAMs (**Fig. 4J**). We then directly compared the pharmacological and both genetic depletion models side by side using an identical 5d depletion paradigm. Following a 2d repopulation period, *Sall1*^CreERT2/wt^:*Csf1r*^fl/fl^ mice exhibited an increased accumulation of CLEC12A^+^F4/80^+^ macrophages in the brain compared to *Cx3cr1*^CreERT2/wt^:*Csf1r*^fl/fl^ and the control condition (**Fig. 4K**).

Altogether, these findings demonstrated that transient niche opening was sufficient for CNS engraftment when CMs are preserved, indicating that the requirement for prolonged niche availability is model-dependent and reflects the concomitant impairment of CM.

### MDMs repopulate CNS niches via clonal expansion, a transient BAM state and direct transpial entry

We next investigated how monocytes access and repopulate CNS compartments. In depleted head-shield chimeras, repopulating macrophages formed discrete TdTomato⁺ clusters in both the brain parenchyma and LM (**Fig. 3D**, **4C, H**). Early repopulation stages revealed isolated seeding events in the LM that subsequently expanded into larger clusters, while columnar arrangements of labeled cells extending from the leptomeningeal surface into the cortex suggested directional infiltration from CNS interfaces (**Fig. 3D**, **4C**, **H**). To directly assess clonal expansion, we generated dual-reporter head-shield chimeras using mixed TdTomato- and GFP-labeled donor BM. This approach enables the assessment of clonal proliferation by distinguishing three distinct MDM sources: donor 1 (TdTomato^+^GFP^-^), donor 2 (TdTomato^-^GFP^+^), and recipient-derived (TdTomato^-^GFP^-^) macrophages (**Fig. 5A**). Following repopulation, MDMs in the LM and parenchyma formed discrete, minimally intermixed donor-derived clusters, demonstrating significant clonal proliferation by Monte Carlo analysis (**Fig. 5A–D**). In contrast, the dura and CP showed substantially greater intermixing and weaker spatial segregation (**Fig. S4G–H**).

**Figure 5:**
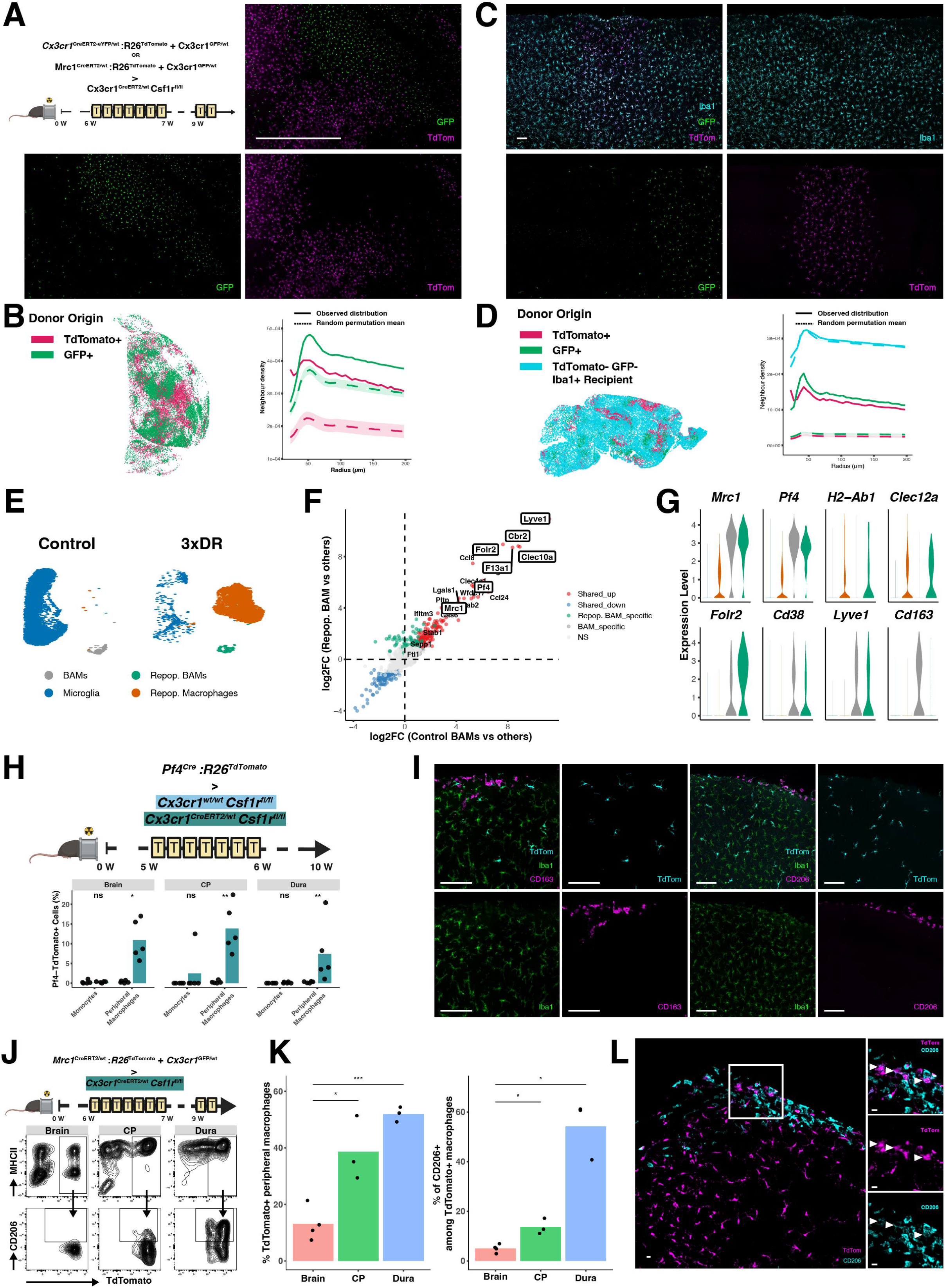
MDMs repopulate CNS niches via clonal expansion, a transient BAM state and direct transpial entry. (**A**) Experimental design and representative en face confocal image of the dorsal brain surface used for spatial analysis. Scale bar: 1mm. (**B**) Left, representative spatial dot plot derived from dorsal en face confocal imaging of the LM. Monte Carlo spatial permutation analysis comparing observed MDM distributions with distributions expected from random permutation of cell identities. Shaded regions indicate the simulation envelope. n=8 (4 Mrc1, 4 Cx3cr1) (**C**) Representative image of spatial distribution of GFP^+^ or TdTomato^+^ Iba1^+^ repopulating MDMs in brain sections from (A). Scale bar: 100µm (**D**) Left, representative dot plot of a sagittal brain section depicting the spatial distribution of tdTomato□, GFP□, and reporter^-^ recipient-derived macrophages. Right, as in (B) but with additional class for Iba1^+^GFP^-^TdTom^-^. n=10 mice (6 Mrc1, 4 Cx3cr1), performed across the brain parenchyma, with an additional class for Iba1^+^GFP^-^TdTom^-^. (**E**) UMAP visualization of published 3xDR scRNA-seq data^49^ highlighting contrasts. (**F**) Differentially expressed genes of control and repopulating BAMs. The upper right quadrant highlights BAM-associated genes re-induced in repopulating BAMs. Log2FC > 0, adj. P.Value > 0.05, pct.1 > 0.25. (**G**) Violin plots of canonical BAM genes in clusters in (E). (**H**) Experimental design and quantification of Pf4-TdTomato⁺ monocytes and MDMs across tissues. See Fig. S4M for gating strategy. n=5 mice per group; one experiment. (**I**) Representative brain sections from (H) showing TdTomato□IBA1□ parenchymal macrophages lacking CD163 and CD206 expression. Scale bar: 100µm. See **Fig. S4L** for TdTomato^+^ leptomeningeal repopulating BAMs. (**J**) Experimental design and representative gating of TdTomato⁺ repopulating macrophages and CD206 expression across tissues. (**K**) Frequency of TdTomato⁺ peripheral macrophages and CD206⁺ cells among TdTomato⁺ peripheral macrophages. (**L**) Representative brain section showing CD206⁺TdTomato⁺ leptomeningeal macrophages and CD206⁻TdTomato⁺ repopulating parenchymal macrophages. Scale bar, 10μm. Statistical tests: (F): Wilcoxon rank-sum test (logfc.threshold = 0, min.pct = 0.1, p.adj < 0.05, genes with pct.1 > 0.25 were displayed). (H, K) Unpaired two-sided Welch’s t-test.

We consistently observed that regions enriched for leptomeningeal TdTomato⁺ macrophages spatially aligned with cortical areas containing parenchymal TdTomato⁺ cells. To visualize this relationship, we performed light-sheet imaging of a repopulated brain hemisphere at d17 (**see Fig. 4H**), which revealed a concordant distribution of leptomeningeal and parenchymal TdTomato⁺ macrophages across large tissue volumes (**Video S1**).

We reasoned that brain-engrafting monocytes would transiently acquire a BAM-like intermediate state during their sequential transition from the blood through the LM and into the brain parenchyma. To identify markers/reporter mice suitable for tracing such a BAM-intermediate state, we leveraged the scRNAseq dataset from the 3xDR model^49^. We specifically searched for markers that were (i) expressed in mature BAMs, (ii) absent from circulating monocytes, and (iii) induced in repopulating leptomeningeal MDMs. We identified *Mrc1* (CD206) and *Pf4* as candidate markers (**Fig. 5E-G**). Another prototypical BAM marker, *CD163*, was excluded because it was neither transcriptionally expressed in monocyte-derived repopulating BAMs (**Fig. 5G** and **Fig. S4I**) nor detected in TdTomato⁺ repopulating leptomeningeal BAMs at d7 or d17 of repopulation in *Cx3cr1*^CreERT2/wt^:*Csf1r*^fl/fl^ head-shield chimeras (**Fig. S4J,** experimental detail in **Fig. 4A, F**). A prerequisite for tracing BAM-intermediate states is the absence of reporter activation in CM. Accordingly, we assessed TdTomato expression following 5d tamoxifen treatment and detected no recombination in *Mrc1*^CreERT2/wt^:*R26*^TdTomato^ and *Lyve1*^CreERT2/wt^:*R26*^TdTomato^ and only minimal (<1.5 %) recombination in *Pf4*^Cre/wt^:*R26*^TdTomato^ mice (**Fig. S4K**).

Next, we performed state-mapping using *Pf4*^Cre/wt^:*R26*^TdTomato^ BM transplanted into *Cx3cr1*^CreERT2/wt^:*Csf1r*^fl/fl^ recipients. Following depletion and repopulation, *Pf4*-TdTomato⁺ macrophages, were detected across all CNS compartments, including the LM (**Fig. S4L**), CP, dura, and brain parenchyma whereas monocytes in the respective tissues remained TdTomato⁻ **(Fig. 5H-I**, see gating strategy in **Fig. S4M)**. These findings indicate that Pf4 expression is induced by monocytes after tissue entry and during their differentiation into BAMs.

*Mrc1* is reported to be rapidly downregulated by brain-engrafting progenitors during the embryonic brain colonization^10,51^. To capture the transient nature of the BAM-state during LM transition, we next generated head-shield chimeras using *Mrc1*^CreERT2/wt^:*R26*^TdTomato^ BM cells (mixed 1:1 with *Cx3cr1*^GFP/wt^) transplanted into *Cx3cr1*^CreERT2/wt^:*Csf1r*^fl/fl^ recipients (**Fig. 5J**). Under this model, monocytes adopting a BAM-like state at the CNS borders would be identified as TdTomato⁺CD206⁺ cells, whereas cells subsequently entering the parenchyma would be expected to downregulate CD206 while retaining permanent TdTomato labeling. Thus, the presence of TdTomato⁺CD206⁻ parenchymal macrophages would indicate that brain-engrafting monocytes transiently passed through a BAM-like state at the LM interface before entering the CNS parenchyma. HDFC analysis revealed that a substantial fraction of macrophages in the dura and CP was TdTomato⁺, whereas recombination was markedly lower in peripheral macrophages in the brain parenchyma (**Fig. 5J-K**). Importantly, within the TdTomato⁺ populations, CD206 protein expression was retained in border compartments but almost completely absent in the brain consistent with local niche imprinting driving CD206 induction and repression, respectively (**Fig. 5K**).

Histological analysis confirmed the presence of CD206⁺TdTomato⁺ leptomeningeal macrophages, whereas brain-engrafting TdTomato⁺ MDMs downregulated CD206 expression (**Fig. 5L**). Collectively, these findings demonstrated that monocytes enter at CNS interfaces, transiently acquire a BAM state and subsequently seed the underlying parenchyma via transpial migration.

### BAMs are a local reservoir for brain parenchymal macrophages

In consequence, we investigated whether MHCII^low^ BAMs themselves, when spared from depletion, could serve as a local source of parenchymal repopulation.

To test this hypothesis, we employed the *Sall1*^CreERT2/wt^:*Csf1r*^fl/fl^ model, which enables selective ablation of MG while preserving CM and BAM populations (**Fig. 4I-J and Fig. S4E-F**). We focused on early repopulation (d7) to capture transitional states prior to phenotypic adaptation to the brain-specific environment^51^. HDFC revealed a marked accumulation of CLEC12A*^+^*F4/80^+^ peripheral macrophages within the brain (**Fig.6A and Fig. S5A**). Within this population, we observed an enrichment of BAM-associated features, including the expansion of MHCII^high^, double negative as well as FOLR2^int^CD206^low^ macrophages, consistent with a transitional differentiation state of BAMs downregulating CD206 (**Fig. 6B**).

**Figure 6:**
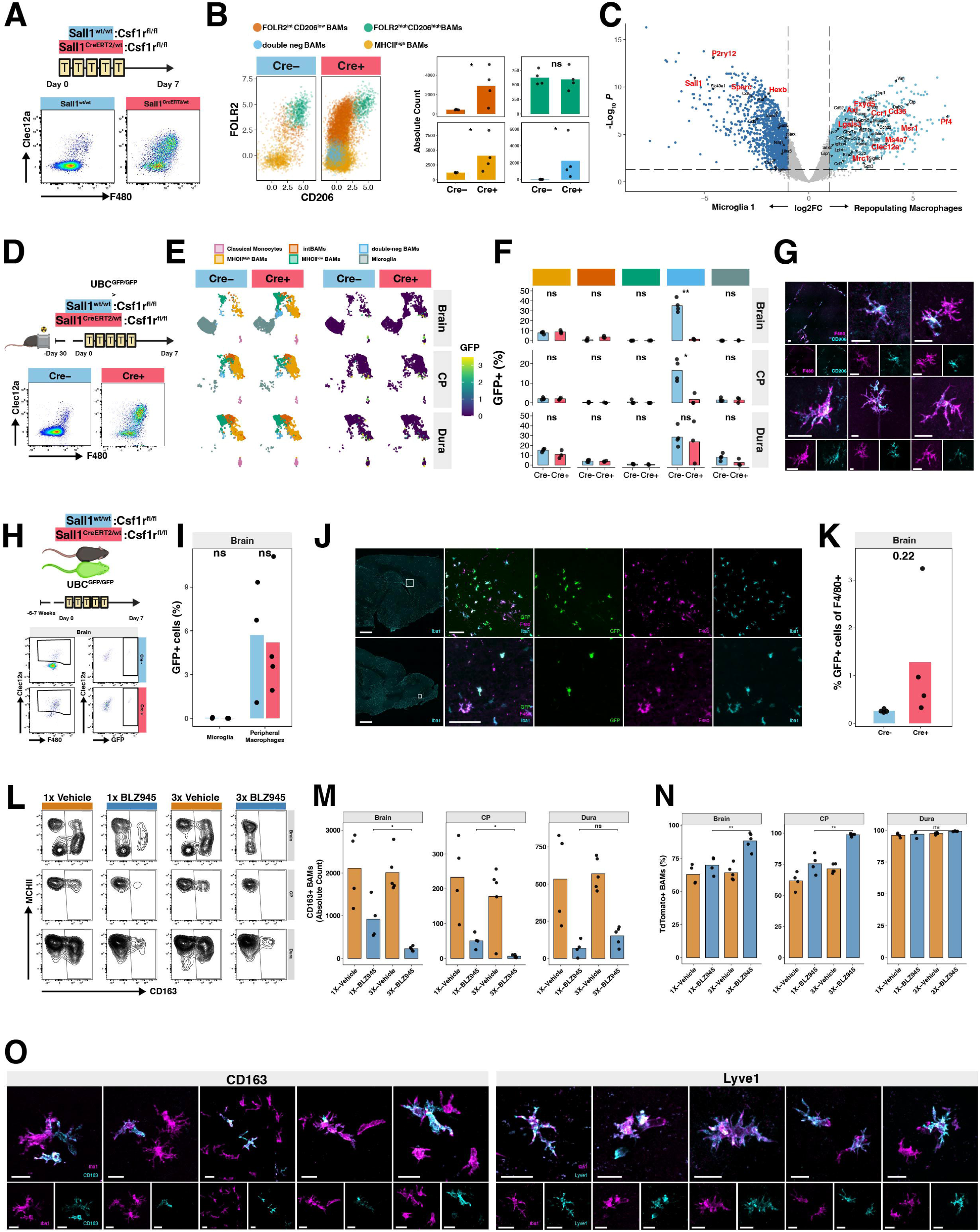
BAMs are a local reservoir for brain parenchymal macrophages. (**A**) Top, experimental scheme and representative FC plots of pre-gated (CD45^+^CD11b^+^Ly6C^-^CD64^+^) macrophages used for cluster analysis in Fig. S5A. (**B**) Left, FC plot of Clec12a+ macrophages from (A). Right, absolute counts of macrophage subsets from (B) (**C**) Volcano plot of differentially expressed genes between Microglia_01 and Repopulating macrophages (see **S5C** for clusters) from scRNAseq experiment in Fig. S5B in tumor-free hemispheres. (**D**) Experimental scheme and representative FC plot showing CD45^+^CD11b^+^Ly6C^-^CD64^+^ pre-gated macrophages used for clustering analysis (**E**) Annotated UMAP and GFP expression overlay of CNS macrophages (Ly6C^+^CCR2^+^ CM were also included into the analysis) from mice in (D). (**F**) Percentage of GFP^+^ cells in clusters shown in (E) n=4 Cre-, n=3 Cre+, one experiment. (**G**) Representative brain sections from (D) showing GFP^-^CD206^+^F480^+^, intraparenchymal macrophages in *Sall1*^CreERT2/wt^*Csf1r*^fl/fl^ repopulated mice (for absence of GFP see **Fig. S6E**. Scale bar: 20 µm. (**H**) Top, experimental outline of parabiosis of *Sall1^CreERT^*^2^*:Csf1r^fl/fl^*and *Sall1^wt/wt^:Csf1r^fl/f^*^l^ mice with UBC^GFP/GFP^ mice (n=3 control, n=4 depletion, from one experiment). Blood chimerism in Fig. S6G. Bottom, representative FC plot showing pre-gated (CD45^+^CD11b^+^Ly6C^-^CD64^+^) macrophages and GFP^+^ cells among Clec12a^+^F480^high^ peripheral macrophages used for quantification. (**I**) Percentage GFP^+^ cells from (**H**). (**J**) Representative brain sections from two different mice from (H) showing rare clusters of GFP^+^Iba1^+^F480^+^ as well as more abundant GFP^-^Iba1^+^F480^+^ intraparenchymal macrophages. Scale bar: 1mm, 100µm. (**K**) Percentage of GFP^+^ cells among F4/80^+^ macrophages quantified by immunofluorescence from (J). (**L-N**) FC analysis of BAMs in brain, CP and dura from experiment outlined in Fig. 3I of repetitive (1xDR, 3xDR and controls) depletion/repopulation using *Flt3^Cre/wt^:R26^TdTomato^*monocyte-reporter mice. (**L**) Representative FC plots of BAMs (pre-gated on F480^high^CD206^+^ macrophages) gating for CD163+ BAMs. (**M**) Absolute counts of CD163+ BAMs as in (L). (**N**) Percentage of TdTomato+ BAMs. (**O)** Representative brain sections from (D) showing GFP-negative CD163^+^Iba1^+^ and Lyve1^+^Iba1^+^ intraparenchymal macrophages in *Sall1*^CreERT2/wt^:*Csf1r*^fl/fl^ repopulated mice. For absence of GFP see **Fig. S6H-I**. Scale bar: 10 µm and 20 µm. Statistical tests: (C, F, I, K, M, N) Unpaired two-sided Welch’s t-test.

However, determining whether BAM-like cells arise from resident BAMs or represent newly recruited monocytes transitioning through a BAM intermediate state remains challenging. To resolve this, we incorporated a tumor-bearing hemisphere as an internal reference, providing a well-defined inflammatory environment that promotes robust monocyte recruitment. Transgenic mice were intracranially injected with syngeneic glioblastoma CT2A tumor cells and, following confirmation of tumor engraftment at d7, treated with tamoxifen for five consecutive days. Tumor-bearing and contralateral tumor-free hemispheres were then collected 7d after depletion onset and processed for scRNAseq (**Fig. S5B**).

Clustering and UMAP embeddings displayed a loss of homeostatic MG (Microglia 1) in Cre^+^ mice with simultaneous appearance of a repopulating macrophages cluster projecting between tumor-associated macrophages and MG (**Fig. S5C**). Subsequent analyses were restricted to the non-tumor-bearing hemisphere to minimize confounding effects driven by inflammation.

Differential gene expression analysis comparing repopulating macrophages to homeostatic MG revealed a strong enrichment of BAM genes, including *Pf4*, *Msr1*, *Mrc1*, *Cd36*, *Ms4a7*, *Clec12a*, and *Axl*, accompanied by reduced expression of canonical MG markers such as *P2ry12*, *Sall1*, *Sparc* and *Hexb* in repopulating macrophages (**Fig 6C**).

To further characterize repopulating macrophages, we reclustered this population and identified three principal populations. The major population exhibited a BAM-like transcriptional profile, defined by high expression of *Pf4*, *Mrc1*, *Clec12a*, *Cxcl16*, and *Ccr1* genes. A second, smaller cluster displayed features of disease-inflammatory macrophages (DIM-like; *Nfkbiz*, *Ccl4*, *Cd14*, *Il1b*, *Cd83*, *Tnfa*), while a third population exhibited a MG-like phenotype, expressing genes such as *Sparc, Hexb, Siglech, and Olfml3.* In addition, each cluster contained a proliferating subpopulation, indicating active self-renewal and expansion during repopulation (**Fig. S5D**).

We validated our BAM-like annotation by projecting the gene signature onto the Macrophage-Verse^6^, a murine reference atlas for CNS macrophages across developmental and adult stages, where the signature exhibited maximal enrichment within annotated BAM populations (**Fig. S5E**).

### Repopulating macrophages do not derive from microglia

To exclude dedifferentiated microglia as a source of the peripheral macrophages, we performed lineage tracing using *Sall1*^CreERT2/wt^:*Csf1r*^fl/fl^:*R26*^TdTomato^ mice (**Fig. S5F**). Repopulating macrophages at d7 were largely TdTomato⁻, with only a minor fraction displaying low-level TdTomato-signal consistent with phagocytic uptake of fluorescent material from dying MG (**Fig. S5G**). These data demonstrated that repopulating macrophages arose independently of mature Sall1^+^ MG. We identified *Pf4* among the most upregulated genes in repopulating macrophages (**Fig.6C**). While Pf4 is a prototypical BAM marker it is also expressed in yolk sac progenitors and primitive macrophages during early CNS colonization^52,53^. To exclude a potential CNS endogenous MG progenitor origin of Pf4⁺ cells that could preferentially repopulate the brain, we first assessed the specificity of *Pf4* expression using *Pf4*^Cre/wt^:*R26*^TdTomato^ reporter mice. In this model, recombination robustly labeled BAM populations across CNS interfaces. However, a non-negligible fraction of TdTomato⁺ cells was also detected within P2Y12⁺ MG (**Fig. S5H**). To resolve the identity of these cells, we performed FLASHseq of TdTomato⁺ cells isolated from brain and dura. This analysis revealed two transcriptionally distinct populations (**Fig. S5I**): a BAM cluster, predominantly derived from the dura, expressing *Pf4*, *Mrc1*, and *Cd163*, and a brain-derived MG cluster expressing canonical MG genes, including *Sall1* and *P2ry12* while lacking *Pf4* expression. *Pf4*-TdTomato^+^ MG were further dependent on CSF1R signaling, as demonstrated by their depletion following CSF1R inhibition (**Fig. S5J**). Moreover, in dual *Sall1*^GFP/wt^:*Pf4*^Cre/wt^:*R26^T^*^dTomato^ reporter mice, P2y12⁺*Pf4*-TdTomato⁺ parenchymal cells show active *Sall1* expression by HDFC and immunofluorescence, confirming their identity as bona fide MG rather than an endogenous progenitor population (**Fig. S5K**).

### Repopulating macrophages do not derive from circulating or skull-derived monocytes

Having excluded a MG origin, we next tested whether repopulating macrophages could derive from CM. We generated head-shield chimeras by transplanting *UBC*^GFP/GFP^ donors in *Sall1*^CreERT2/wt^:Csf1r^fl/fl^ and *Sall1*^wt/wt^:*Csf1r*^fl/fl^ control mice (**Fig. 6D**), achieving a ∼20–40% chimerism in CM (**Fig. S6A)**. On d7 after depletion start, GFP^+^ cells were detected within subdural and dural/CP BAM populations across both conditions but did not show increased contribution upon depletion, consistent with baseline monocyte turnover of sdBAMs (**Fig. 6E-F**, see **Fig. S6B** for cluster identity projected on conventional plot (left) and GFP density plot per subset (right) and conventional gating/analysis in **Fig. S6C-D**).

Histological analysis identified only rare GFP⁺F4/80⁺ MDMs with the majority of repopulating parenchymal F4/80⁺ macrophages remaining GFP⁻, while frequently expressing CD206 (**Fig. 6G**, see **Fig. S6E** for GFP channel). Importantly, no differential increase in GFP^+^ macrophage abundance was observed across conditions (**Fig. S6F**).

To further corroborate our findings, we performed parabiosis experiments by surgically conjoining UBC^GFP/GFP^ reporter mice with either *Sall1*^CreERT2/wt^:*Csf1r*^fl/fl^ or *Sall1*^wt/wt^:*Csf1r*^fl/fl^ control mice. After 6–7 weeks, stable blood chimerism of approximately 25% was established (**Fig. S6G**), comparable to head-shield chimeras. Mice were then treated with tamoxifen and analyzed 7d after treatment (**Fig. 6H**). HDFC revealed no enrichment of GFP^+^ cells within the F4/80^high^CLEC12A^+^ repopulating macrophages or within MG across groups (**Fig. 6I**). In line with our head-shield chimeras, GFP^+^ cells were detectable within the peripheral macrophage/BAM gate in non-depleted controls, indicating that they reflect baseline monocyte contribution to sdBAMs rather than depletion-induced recruitment of monocytes into the parenchymal niche (**Fig. 6H-I**).

Overall, parenchymal F480^high^ macrophages were predominantly GFP^-^, with only rare GFP^+^F4/80^high^ parenchymal cells observed, largely restricted to the olfactory bulb or periventricular regions with no significant difference of GFP^+^ cells among F480^high^ macrophages (**Fig. 6J-K**).

Although skull-derived monocytes can engraft a chronically microglia-deficient niche as recently reported^54^, UBC^GFP/GFP^ calvarial bone-flap transplantation onto *Sall1*^CreERT2/wt^:*Csf1r*^fl/fl^ mice revealed no detectable contribution of skull-derived macrophages to the repopulating parenchymal macrophage pool following transient microglia-selective depletion, as assessed by HDFC and immunofluorescence (**Fig. S6J**).

### CD163 identifies BAM-derived macrophages and is not re-induced in MDMs

Next, we sought to identify lineage-restricted markers distinguishing embryonic BAMs from monocyte-derived MHCII^high^ BAMs. For this, we leveraged the 3xDR model (**Fig. 5G** and **Fig. S4I**), in which iterative CSF1Ri progressively exhausted the self-renewal capacity of MG and promoted their replacement by MDMs.

Differential gene expression analysis revealed that *Cd163 was* selectively expressed in embryonic BAMs (in control condition) but not re-induced in repopulating BAMs. In contrast, other MHCII^low^ BAM-associated markers, including *FOLR2, CD38, and Lyve1*^26^, were re-expressed in repopulating BAMs, indicating that these markers reflect tissue adaptation rather than lineage origin (**Fig. 5F-G**).

To validate these findings, we analyzed BAMs in the repetitive depletion/repopulation model (**Fig.3I**). CD163⁺ BAMs progressively declined with each successive depletion round, most prominently within the subdural and CP niches, while the overall fraction of TdTomato⁺ BAMs increased, consistent with gradual replacement by CD163⁻ MDMs (**Fig. 6L-N**). Taken together, these data demonstrate that CD163 is not induced upon tissue entry, within the examined time frame, establishing it as a lineage-restricted marker of bona-fide mature BAMs.

We stained brains of *Sall1*^CreERT2/wt^:*Csf1r*^fl/fl^ head-shield chimeras (**Fig. 6D**) for CD163 and LYVE1 to determine whether BAMs could be detected within the repopulated brain parenchyma. Indeed, IBA1⁺CD163⁺GFP^⁻^ and IBA1⁺LYVE1⁺GFP^⁻^ macrophages were readily observed within the parenchyma (**Fig. 6O**). Given the short time they resided in this compartment, the presence of CD163⁺/Lyve1^+^ macrophages strongly supports retention of a pre-existing lineage identity rather than de novo induction. Together, these data show that BAMs can be a local reservoir for brain parenchymal macrophages.

### Peripheral macrophages are detectable in the aged human brain independent of Alzheimer’s pathology

Given that chronic proliferative stress may progressively exhaust long-term MG self-renewal, we asked whether peripheral macrophages can be detected within the aging human CNS parenchyma when MG are no longer able to sustain local niche demand under physiological or neurodegenerative conditions. We leveraged a scRNAseq brain atlas of aged human individuals with and without Alzheimer’s disease (AD) from the Religious Orders Study and Memory and Aging Project (ROSMAP)^55^. The dataset comprises samples from six brain regions: the entorhinal cortex (EC), hippocampus (HC), anterior thalamus (TH), angular gyrus (AG), middle temporal cortex (MT), and prefrontal cortex (PFC). Within the human brain macrophage landscape, we identified two clusters (C6 and C12) strongly enriched for the core murine MDM signature (**Fig. 7B**). Both clusters expressed CD163; while a further cluster C10 was enriched for a published human BAM signature^56^, consistent with a bona fide BAM identity (**Fig. 7C**). In contrast, C6 was further characterized by elevated expression of peripheral macrophage markers (ITGA4, LGALS3, CXCR4, FXYD5, C5AR1), together with disease-associated microglia (DAM)-related genes (GPNMB, ITGAX, TREM2 and CD9) and BAM-associated genes (CD163, SIGLEC1, F13A1, and MSR1) (Fig. 7D). C6 and C12 were reciprocally enriched in AD and non-AD brains respectively (**Fig. 7A**) while being consistently detected across all brain regions (**Fig. 7E**). To localize these cells within the human brain parenchyma, we performed immunofluorescence staining on postmortem human brain sections obtained from AD patients (n=2, including one individual with trisomy 21-associated AD pathology) as well as one age-matched non-AD (99 years old) control (**Fig. 7A**). Analyses were conducted across multiple anatomical brain regions. We focused on CD163 as a lineage-associated peripheral macrophage marker, in combination with IBA1 to identify CNS-resident macrophages. Consistent with recent reports that CD163 marks human MDMs^26,56,57^, we detected ramified CD163⁺IBA1⁺ cells within the brain parenchyma across multiple anatomical regions in both control and disease samples. Notably, CD163⁺IBA1⁺ parenchymal cells were detected in relatively AD-spared regions such as the substantia nigra of AD patients as well as in multiple regions of an aged non-AD individual, including the calcarine sulcus and medulla oblongata (**Fig. 7I**). Their presence across multiple brain regions and conditions suggests that peripheral macrophages colonize the human CNS parenchyma under physiological and pathological states.

**Figure 7:**
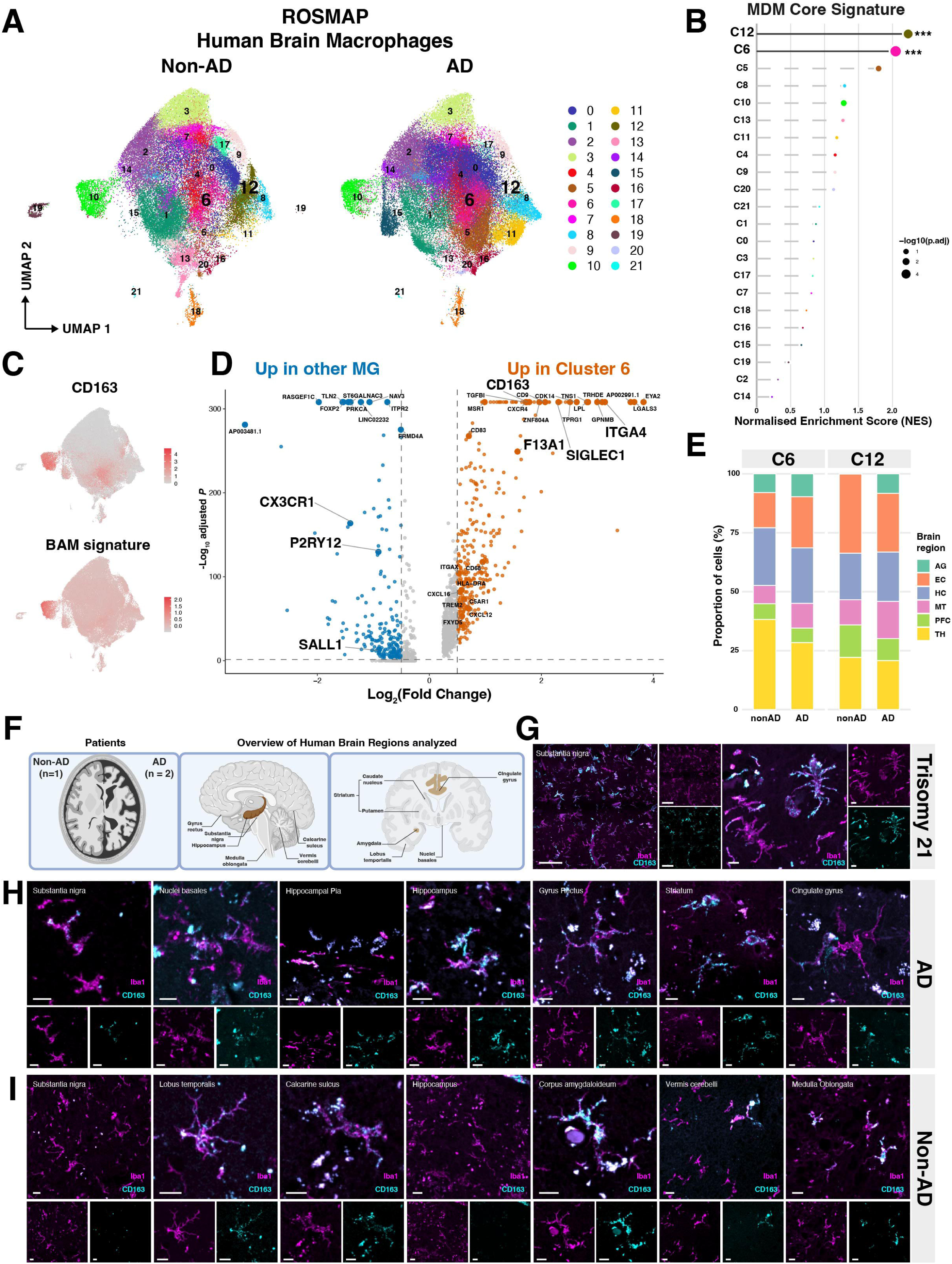
Peripheral macrophages are detectable in the aged human brain independent of Alzheimer’s pathology. (**A**) UMAP of scRNAseq human brain macrophages (ROSMAP^55^). (**B**) Enrichment analysis of murine core MDM signature onto human CNS macrophages. (**C**) Feature plot of CD163 and AddModuleScoring of published^56^ human BAM signature. (**D**) Differentially expressed genes between C12 and MG. (**E**) Regional composition of C6/C12. (**F**) Overview of human postmortem brain samples analyzed by immunofluorescence imaging. (**G-I**) Representative brain sections of (**G**) trisomy 21, (**H**) AD patient and (**I**) aged control. Scale bar : 100 µm, 10 µm.

Together, these findings suggest that the aging human brain macrophage landscape contains a previously underappreciated fraction of non-MG peripheral macrophages.

## Discussion

The principles that determine when and how monocytes or peripheral macrophages gain access to CNS compartments remain incompletely understood. Experimental macrophage depletion paradigms have become important tools to investigate CNS macrophage maintenance and function. However, apparently conflicting repopulation outcomes across macrophage depletion strategies^7,15–17^ have complicated interpretation of these studies and contributed to the view that prolonged niche availability dictates peripheral engraftment^26^. Here, we integrated pharmacological and genetic macrophage depletion models to identify shared physiological variables that govern repopulation outcomes across major CNS macrophage niches. We show that these models are not contradictory; rather, they represent distinct perturbations of a common biological framework centered on two key parameters and their temporal alignment: presence of engraftment-competent cells and TRM self-renewal capacity.

Broad pharmacological CSF1Ri efficiently depletes MG but also transiently impairs CM differentiation and engraftment, despite preserving their abundance. Rapidly self-renewing MG re-establish niche dominance and limit peripheral engraftment. In contrast, persistent impairment of TRM self-renewal, through either irreversible genetic disruption of CSF1R signaling or iterative depletion-repopulation cycles, exhausted yolk sac-derived TRMs (including MG, sd- and cpBAMs) and shifted the competitive balance toward peripheral engraftment. Importantly, selective TRM depletion demonstrated that MDM engraftment does not require prolonged niche vacancy if engraftment-competent cells are preserved during transient niche opening. Despite partial acquisition of canonical MG genes (P2y12, Tmem119), brain-engrafting monocytes failed to fully adopt a bona fide MG identity (*Sall1*^27,58^), potentially reflecting ontogenetically imprinted epigenetic constraints^26,54^ or insufficient time within the CNS niche^59^. We show that brain-engrafting MDMs are transcriptionally heterogeneous yet share a conserved core MDM transcriptional program, which, consistent with recent reports^54,60^, includes acquisition of a BAM-like transcriptional profile. We extended these findings by mapping the spatiotemporal acquisition of the BAM state by CNS-infiltrating monocytes, demonstrating that monocytes progressively differentiate through a BAM identity as they propagate from blood to the LM and, under permissive conditions, into the adjacent brain parenchyma. Consistent with the transpial seeding of the macrophage-devoid embryonic brain^54^ and the postnatal colonization of the perivascular space by leptomeningeal BAMs^61^, we show that a re-opened parenchymal niche in adult mice is again accessible directly through the pial surface to monocyte-derived MHCII^high^ and embryonic MHCII^low^ CD206^+^/Lyve1^+^/CD163^+^ BAMs. Together, these findings suggest that the LM serve not solely as a barrier, but as a macrophage-brain entry interface.

We found that the subdural compartment and the brain parenchyma constitute a restrictive niche in which monocyte entry is limited and dependent on Ccr2. In contrast, the dura and CP support continuous monocyte influx and undergo replenishment independently of Ccr2, consistent with recent studies^11,13,26,62^ and closely mirroring replacement dynamics observed in human HSC-transplanted patients^56^. Importantly, these accessibility gradients dictate distinct modes of niche reconstitution. In restricted compartments repopulation is dominated by local clonal expansion following sporadic seeding events, as limited influx provides newly engrafted cells with a prolonged window for local proliferation before encountering competing cells.

Peripheral macrophage engraftment into the brain parenchyma has been documented in aging mice^59^, in neurodegeneration in both mice^63,64^ and humans^26,65^, as well as in HSC-transplanted leukemia patients^56^. Although definitive lineage tracing of brain-engrafting MDMs remains largely restricted to mice, converging human evidence supports their presence in the parenchyma. Sex-mismatched HSC transplantation studies identified Y-chromosome-positive donor-derived macrophages and, by single-nucleus RNA sequencing, an enrichment of CD163-expressing macrophages in transplanted brains^56^, while xenotransplantation studies independently supported CD163 as a marker of brain-engrafting human MDMs^26,57^. Together with reports of CD163+ macrophage accumulation in perivascular and subpial regions during neurodegeneration^65^, these findings suggest that peripheral macrophages can access the human parenchyma through routes consistent with transpial entry. Our data broaden the evidence for CD163⁺IBA1⁺ parenchymal macrophages by demonstrating their presence across multiple brain regions in both aged and neurodegenerative human brains, including individuals without overt neuropathology.

This interpretation is further supported by a recent spatial proteomic analysis of human AD and aged control brains, which similarly identified a BAM-like MG population co-expressing CD163 and TMEM119^66^, likely reflecting a population analogous to the population observed in our study.

Notably, both detrimental and beneficial effects of brain-engrafting monocytes have been reported in mouse^49,54,57,59^ and human^67,68^, indicating that their functional impact is not uniform. It therefore remains to be determined which mechanisms govern these divergent outcomes, potentially including origin, route of engraftment and pre-conditioning prior to CNS entry, as well as the local niche context that shapes their functional integration.

The ability to predictably repopulate even the highly restrictive microglial niche suggests that similar principles govern macrophage niche occupation across tissues beyond the CNS. We provide a new conceptual framework for the design of macrophage replacement strategies, which have historically been complicated by the adverse effects associated with irradiation- or chemotherapy-based conditioning^69^. In this context, irradiation- and chemo-free approaches based on iterative TRM exhaustion already represent a significant advance as recently suggested^54^. We show that coupling controlled niche opening with the preservation of engraftment-competent cells or, alternatively, with the introduction of shielded exogenous cells enables rapid, directed, and controlled repopulation. This strategy minimizes the duration of niche vulnerability while improving temporal control and cellular composition of niche reconstitution.

Our findings further reveal that CNS border compartments are not impenetrable barriers, but conditionally permissive interfaces through which macrophages can transition across anatomically connected niches. In this framework, the LM emerge as an immunologically active reservoir and conditional gateway for macrophage entry into the brain parenchyma.

## Limitations of the study

While we demonstrate that monocyte-derived MHCII^high^ BAMs undergo transpial entry into the brain parenchyma, selective genetic fate-mapping strategies for embryonic MHCII^low^ BAMs are currently unavailable. Consequently, because of the remarkable plasticity and rapid phenotypic convergence of macrophages during niche adaptation, the exact relative contribution of MHCII^high^ and MHCII^low^ BAMs to microglial replacement cannot yet be definitively quantified. Furthermore, whether similar permissive niche conditions and transpial recruitment of BAMs occur during physiological aging or distinct pathological conditions, rather than acute experimental niche perturbation, remains to be determined.

Finally, although our transcriptomic and histological analyses support the presence of peripheral macrophages within the human brain, definitive lineage relationships cannot currently be established in humans. Future studies integrating naturally occurring somatic lineage markers, including mitochondrial DNA variants and clonal hematopoiesis associated mutations, with single-cell multiomic and spatial analyses may enable direct reconstruction of macrophage ontogeny and replacement dynamics in humans.

## Supporting information

Graphical Abstract

## RESOURCE AVAILABILITY

### Lead contact

- Requests for further information and resources should be directed to and will be fulfilled by the lead contact, Gregor Hutter (^29,30^).

### Materials availability

This study did not generate new unique reagents.

### Data and code availability

All single-cell RNA-seq datasets generated in this study will be deposited in the Gene Expression Omnibus (GEO) and made publicly accessible under the corresponding GEO accession code upon publication. Additional processed data and code required to reproduce the analyses is available from the lead contact upon request.

SnRNA-seq profiling data are available from Synapse in coordination with the ROSMAP project. Data are accessible under accession codes syn52293442 (as part of the MIT ROSMAP Single-Nucleus Multiomics Study; Synapse: syn52293417). The data are available under controlled use conditions set by human privacy regulations. To access the data, a data use agreement is needed. This registration is in place solely to ensure anonymity of the ROSMAP study participants. A data use agreement can be agreed with either Rush University Medical Center (RUMC) or with SAGE, who maintains Synapse, and can be downloaded from their websites (https://www.radc.rush.edu/; https://adknowledgeportal.synapse.org/).

## ACKNOWLEDGMENTS

Calculations were performed at sciCORE (http://scicore.unibas.ch/) scientific computing center at the University of Basel. This work was supported by a Swiss National Science Foundation Research MD-PhD Grant (323630-214538) to D.K.; Swiss National Science Foundation Professorial Fellowship (PP00P3_176974); Swiss National Science Foundation Project grant (3200-0-239894) to G.H. the ProPatient Forschungsstiftung, University Hospital Basel (Annemarie Karrasch Award 2019); Swiss Cancer Research Grant (KFS- 4382-02-2018) to G.H.; the Department of Surgery, University Hospital Basel, to G.H. and D.K.; We thank Prof. Alfred Zippelius, Prof. Jan Niess and Prof. Ryuichi Nishinakamura for providing genetic mouse lines; the genomics, proteomics, flow cytometry histology, microscopy and animal core facilities of the University of Basel, Switzerland for technical and logistical support. We thank Alexandar Tzankov for providing human brain tissue samples. We thank Diego Calabrese and the members of the DBM Histology Core Facility for their support and staining service. We thank the microscopy facilities at the Department of Biomedicine (DBM), particularly Michael Abanto, and the Imaging Core Facility (IMCF, Biozentrum), especially Kai Schleicher, for their technical support with microscopy and imaging workflows. We also thank Marc Thommen for advice and support regarding tissue clearing protocols. We are extremely grateful to Paul Jordi from Jordi Strahlentechnik (Münchenstein, Basel) for the design and construction of the head-shielding chamber. We further thank Roman Menz (University Hospital Basel) and Thibaut Klein (Department of Biomedicine, Basel) for their support in planning the irradiation experiments. We thank the study participants and staff of the Rush Alzheimer’s Disease Center, the data contributors, and the investigators of the original study. The dataset was originally generated and described by Mathys, H., Boix, C.A., Akay, L.A. et al. Single-cell multiregion dissection of Alzheimer’s disease. Nature 632, 858–868 (2024). https://doi.org/10.1038/s41586-024-07606-7.

## AUTHOR CONTRIBUTIONS

Conceptualization, D.K, R.Gl., G.H.; methodology, D.K, G.H, P.T.W.; investigation, D.K., P.T.W., S.H., G.D.B, S.D., S.Z., M.F.R., T.S., H.M., Z.B., V.S., F.G.,; formal analysis, D.K, P.T.W, S.H., P.Z.,; writing-original draft, D.K; writing-review & editing, D.K., P.T.W., G.D.B., A.C.D., M.F.R., and G.H.; funding acquisition, D.K. and G.H.; supervision, G.H; data curation, D.K, M.F.R, G.D.B., S.Z., R.G., S.D., A.G., M.M.; project administration, M.F.R., G.D.B.; Resources, G.H., M.F.R., A.G., M.D., A.Z., R.G., L.M., J.K., R.Guz.

## DECLARATION OF INTERESTS

G.H. has equity in, and is a co-founder of Incephalo Inc., G.H. is a co-founder of GlioCART GmbH, G.H. has received consultancy honoraria from Servier Inc., Novartis and BB Braun Aeskulap.

## DECLARATION OF GENERATIVE AI AND AI-ASSISTED TECHNOLOGIES

During preparation of this work, the authors used GPT-5.5 for language, structure, clarity, and readability of the manuscript. The authors reviewed and edited all AI-assisted output as needed and take full responsibility for the accuracy, integrity, and content of the published work.

## SUPPLEMENTAL INFORMATION

Figures S1-S6

S1 Video, related to Fig. 4F./Fig. 5.

Primer List

Spplementary_Table_sample_metadata_ROSMAP

Supplementary_Table_HumanSamples_IF

## STAR⍰METHODS

### KEY RESOURCES TABLE

| REAGENT or RESOURCE | SOURCE | IDENTIFIER |
| --- | --- | --- |
| **Antibodies** |  |  |
| Goat monoclonal anti Iba1 | Novus Biologicals | Cat#NB100-1028;<br>RRID: AB_521594 |
| Goat polyclonal anti Iba1 | Abcam | Cat#ab5076;<br>RRID:AB_2224402 |
| Rat monoclonal anti MHC II | Invitrogen | Cat#14-5231-82;<br>RRID: AB_467561 |
| Rat monoclonal anti CD206 | Thermo Scientific | Cat#MA5-16871;<br>RRID: AB_2538349 |
| Rabbit polyclonal anti Iba1 | Fujifilm Wako | Cat#019-19741;<br>RRID: AB_839504 |
| Rabbit monoclonal anti CD206 | Cell Signaling<br>Technology | Cat#24595; RRID:<br>AB_2892682 |
| Rabbit monoclonal anti TMEM119 | Abcam | Cat#ab209064;<br>RRID: AB_2800343 |
| Rabbit monoclonal anti P2Y12 | AnaSpec | Cat#55043A; RRID:<br>AB_2298886 |
| Rat monoclonal anti Lyve1 | Invitrogen | Cat#14-0443-80;<br>RRID: AB_1633414 |
| Rat monoclonal anti F4/80 | Invitrogen | Cat#14-4801-82;<br>RRID: AB_467558 |
| Rabbit monoclonal anti Ki67 | Cell Signaling<br>Technology | CAT#12202S; RRID:<br>AB_2620142 |
| Goat polyclonal anti Axl | R&D Systems | CAT#AF854; RRID:<br>AB_355663 |
| Chicken polyclonal anti GFP | Aves | CAT#GFP-1010;<br>RRID: AB_2307313 |
| Rabbit monoclonal anti CD163 | Abcam | CAT#ab182422;<br>RRID: AB_2753196 |
| Rat monoclonal anti CD163 | BioLegend | CAT#155302; RRID:<br>AB_2734239 |
| Donkey anti goat AF488 | Jackson | Cat#705-545-147; |
|  | ImmunoResearch | RRID: AB_2336933 |
| Donkey anti rabbit AF555 | Jackson<br>ImmunoResearch | Cat#711-565-152;<br>RRID: AB_3095471 |
| Donkey anti chicken AF555 | Jackson<br>ImmunoResearch | Cat#703-165-155;<br>RRID: AB_2340363 |
| Donkey anti chicken AF488 | Invitrogen | Cat#A-11039; RRID:<br>AB_2534096 |
| Donkey anti goat AF647 | Jackson<br>ImmunoResearch | Cat#705-605-147;<br>RRID: AB_2340437 |
| Donkey anti rabbit AF647 | Jackson<br>ImmunoResearch | Cat#711-605-152;<br>RRID: AB_2492288 |
| Donkey anti goat AF750 | Abcam | Cat#ab175745;<br>RRID: AB_2924800 |
| Donkey anti rat AF647 | Jackson<br>ImmunoResearch | Cat#712-605-153;<br>RRID: AB_2340694 |
| Donkey anti rat AF488 | Jackson<br>ImmunoResearch | Cat#712-545-150;<br>RRID: AB_2340683 |
| Donkey anti rabbit AF488 | Jackson<br>ImmunoResearch | Cat#711-545-152;<br>RRID: AB_2313584 |
| Donkey anti goat CY3 | Jackson<br>ImmunoResearch | Cat#705-166-147;<br>RRID: AB_2340413 |
| Donkey anti-goat AF546 | Jackson<br>ImmunoResearch | Cat#705-165-147;<br>RRID:AB_2307351 |
| AF488 anti CD31 | BioLegend | Cat#102514; RRID:<br>AB_2161031 |
| PE anti Lyve1 | Thermo Fisher<br>Scientific | Cat#12-0443-80;<br>RRID: AB_2802178 |
| RB780 anti CD301b | BD Biosciences | Cat#569476, RRID:<br>AB_3685091 |
| APC anti CD11a | BioLegend | Cat#101119; RRID:<br>AB_2562778 |
| BV421 anti CD13 | BD Biosciences | Cat#752980; RRID:<br>AB_2917935 |
| RB670 anti CD13 | BD Biosciences | Cat#770477; RRID:<br>AB_3692347 |
| AF488 anti Axl | R and D Systems | Cat#FAB8541G;<br>RRID: AB_3653411 |
| PE/Cy7 anti CD31 | BioLegend | Cat#102523; RRID:<br>AB_2572181 |
| AF647 anti CD31 | BioLegend | Cat#102516; RRID:<br>AB_2161029 |
| AF647 anti Tim-4 | BioLegend | Cat#130007; RRID:<br>AB_2201842 |
| PE anti I-A/I-E | BioLegend | Cat#107607; RRID:<br>AB_313322 |
| AF647 anti CD192 | BD Biosciences | Cat#568347; RRID:<br>AB_3684201 |
| AF647 anti F4/80 | BD Biosciences | Cat#565854; RRID:<br>AB_2744474 |
| FITC anti CX3CR1 | BioLegend | Cat#149020; RRID:<br>AB_2565703 |
| BV605 anti CD45 | BioLegend | Cat#103140; RRID:<br>AB_2562342 |
| APC anti CD9 | BioLegend | Cat#124812; RRID:<br>AB_2783071 |
| AF488 anti I-A/I-E | BioLegend | Cat#107616; RRID:<br>AB_493523 |
| RB613 anti CD11c | BD Biosciences | Cat#758187; RRID:<br>AB_3690336 |
| BUV395 anti CD11b | BD Biosciences | Cat#563553; RRID:<br>AB_2738276 |
| PE-CF594 anti CD192 | BD Biosciences | Cat#568804; RRID:<br>AB_3684560 |
| APC/Fire 810 anti P2RY12 | BioLegend | Cat#848014; RRID:<br>AB_2904450 |
| PE/Cy5 anti CD64 | BioLegend | Cat#139331; RRID:<br>AB_2922467 |
| BUV805 anti F4/80 | BD Biosciences | Cat#749282; RRID:<br>AB_2873657 |
| BUV563 anti CD49d | BD Biosciences | Cat#741243; RRID: |
|  |  | AB_2870794 |
| AF700 anti CD11c | BioLegend | Cat#117320; RRID:<br>AB_528736 |
| PE/Cy7 anti CD3ε | BioLegend | Cat#100320; RRID:<br>AB_312685 |
| BUV563 anti Ly-6G | BD Biosciences | Cat#612921; RRID:<br>AB_2870206 |
| PE/Fire700 anti NK-1.1 | BioLegend | Cat#156527; RRID:<br>AB_2910320 |
| APC/Cy7 anti CD19 | BioLegend | Cat#115529; RRID:<br>AB_830706 |
| FITC anti CD9 | BioLegend | Cat# 124807; RRID:<br>AB_1279324 |
| BUV737 anti CD45 | BD Biosciences | Cat#568344; RRID:<br>AB_3684199 |
| PE anti CX3CR1 | BioLegend | Cat#149005; RRID:<br>AB_2564314 |
| APC/Cy7 anti CD86 | BioLegend | Cat#105030; RRID:<br>AB_2244452 |
| BV605 anti CX3CR1 | BioLegend | Cat#149027; RRID:<br>AB_2565937 |
| PE anti CD319 | BioLegend | Cat#152006; RRID:<br>AB_2632677 |
| AF700 anti CD43 | BioLegend | Cat#143214; RRID:<br>AB_2800661 |
| APC anti CD319 | BioLegend | Cat#152003; RRID:<br>AB_2632674 |
| APC/Fire 810 anti CX3CR1 | BioLegend | Cat#149053; RRID:<br>AB_2910300 |
| PE/Cy7 anti CD62L | BioLegend | Cat#104417; RRID:<br>AB_313102 |
| RB780 anti CD115 | BD Biosciences | Cat#755874; RRID:<br>AB_3688171 |
| AF488 anti I-A/I-E | BioLegend | Cat#107615; RRID:<br>AB_493524 |
| AF700 anti CD206 | BioLegend | Cat#141734; RRID:<br>AB_2629637 |
| APC anti CD371 | BioLegend | Cat#143405; RRID:<br>AB_2564264 |
| TruStain FcX | BioLegend | Cat#101320; RRID:<br>AB_1574975 |
| RB613 anti Folate Receptor $\beta$ | BD Biosciences | Cat#759707; RRID:<br>AB_3691779 |
| APC/Fire 810 anti CD163 | BioLegend | Cat#155321; RRID:<br>AB_2904294 |
| BV785 anti Ly6C | BioLegend | Cat#128041; RRID:<br>AB_2565852 |
| PE/Cy7 anti CD38 | BioLegend | Cat#102718; RRID:<br>AB_2275531 |
| FITC anti CD64 | BioLegend | Cat#139316; RRID:<br>AB_2566556 |
| BB700 anti I-A/I-E | BD Biosciences | Cat#746197; RRID:<br>AB_2743544 |
| BV421 anti CD38 | BioLegend | Cat#102732; RRID:<br>AB_2734153 |
| PE anti 371 | BioLegend | Cat#143403; RRID:<br>AB_11126747 |
| Spark Blue 574 anti CD8a | BioLegend | Cat#100793; RRID:<br>AB_2922450 |
| BUV496 anti CD4 | BD Biosciences | Cat#612952; RRID:<br>AB_2813886 |
| BV650 anti CD8a | BioLegend | Cat#100742; RRID:<br>AB_2563056 |
| PE/Cy7 anti CD45.2 | BioLegend | Cat#109829; RRID:<br>AB_1186103 |
| FITC anti CD8a | BioLegend | Cat#100706; RRID:<br>AB_312745 |
| APC/Cy7 anti CD3ε | BioLegend | Cat#100330; RRID:<br>AB_1877170 |
| PE anti CD115 | BD Biosciences | Cat#566839; RRID: |
|  |  | AB_2869896 |
| APC anti Folate Receptor $\beta$ | BioLegend | Cat#153305; RRID:<br>AB_2721312 |
| FITC anti CD9 | BioLegend | Cat#124807; RRID:<br>AB_1279324 |
| APC/Cy7 anti CD19 | BioLegend | Cat#115529; RRID:<br>AB_830706 |
| PE/Fire 700 anti NK1.1 | BioLegend | Cat#156527; RRID:<br>AB_2910320 |
| BUV563 anti Ly-6G | BD Biosciences | Cat#612921; RRID:<br>AB_2870206 |
| PE/Cy7 anti CD3ε | BioLegend | Cat#100320; RRID:<br>AB_312685 |
| AF700 anti CD11c | BioLegend | Cat#117320; RRID:<br>AB_528736 |
| BUV563 anti CD49d | BD Biosciences | Cat#741243; RRID:<br>AB_2870794 |
| BUV805 anti F4/80 | BD Biosciences | Cat#749282; RRID:<br>AB_2873657 |
| PE/Cy5 anti CD64 | BioLegend | Cat#139331; RRID:<br>AB_2922467 |
| APC/Fire 810 anti P2RY12 | BioLegend | Cat# 848014; RRID:<br>AB_2904450 |
| PE-CF594 anti CD192 | BD Biosciences | Cat#568804; RRID:<br>AB_3684560 |
| BUV395 anti CD11b | BD Biosciences | Cat#563553; RRID:<br>AB_2738276 |
| RB613 anti CD11c | BD Biosciences | Cat#758187; RRID:<br>AB_3690336 |
| AF488 anti I-A/I-E | BioLegend | Cat#107616; RRID:<br>AB_493523 |
| APC anti CD9 | BioLegend | Cat#124812; RRID:<br>AB_2783071 |
| BV605 anti CD45 | BioLegend | Cat#103140; RRID:<br>AB_2562342 |
| FITC anti CX3CR1 | BioLegend | Cat#149020; RRID:<br>AB_2565703 |
| AF647 anti CD192 | BD Biosciences | Cat#568347; RRID:<br>AB_3684201 |
| PE anti I-A/I-E | BioLegend | Cat#107607; RRID:<br>AB_313322 |
| AF647 anti Tim-4 | BioLegend | Cat#130007; RRID:<br>AB_2201842 |
| AF647 anti CD31 | BioLegend | Cat#102516; RRID:<br>AB_2161029 |
| PE/Cy7 anti CD31 | BioLegend | Cat#102523; RRID:<br>AB_2572181 |
| AF488 anti Axl | R and D Systems | Cat#FAB8541G;<br>RRID: AB_3653411 |
| RB670 anti CD13 | BD Biosciences | Cat#770477; RRID:<br>AB_3692347 |
| BV421 anti CD13 | BD Biosciences | Cat#752980; RRID:<br>AB_2917935 |
| APC anti CD11a | BioLegend | Cat#101119; RRID:<br>AB_2562778 |
| RB780 anti CD301b | BD Biosciences | Cat#569476; RRID:<br>AB_3685091 |
| PE anti Lyve1 | Thermo Fisher<br>Scientific | Cat#12-0443-80;<br>RRID: AB_2802178 |
| AF488 anti CD31 | BioLegend | Cat#102514; RRID:<br>AB_2161031 |
| Bacterial and virus strains |  |  |
| Biological samples |  |  |
| Chemicals, peptides, and recombinant proteins |  |  |
| DAPI | BD Pharmingen | CAT#564907 |
| PE anti Streptavidin | BioLegend | CAT#405203 |
| Tamoxifen | MedChem Express | CAT#HY-13757A |
| Corn oil | Merck | CAT#C8627 |
| Brilliant stain buffer | Thermo Scientific | CAT#00-4409-75 |
| Collagenase, Type 4 | Worthington | CAT#LS004188 |
|  | Biochemical |  |
| Debris Removal Solution | Miltenyi Biotec | CAT#120-109-398 |
| Collagenase, Type 1 | Worthington<br>Biochemical | CAT#LS004196 |
| DNase I | Millipore Sigma | CAT#10104159001 |
| Cyto-Fast Fix/Perm Buffer Set | BioLegend | CAT#426803 |
| RedSafe Nucleic Acid Staining Solution | INtRON Biotechnology<br>Inc | CAT#21141 |
| Monocyte Isolation Kit (BM), mouse | Miltenyi Biotec | CAT#130-100-629 |
| True-Stain Monocyte Blocker | BioLegend | CAT#426102 |
| AutoMACS Running Buffer | Miltenyi Biotec | CAT#130-091-221 |
| Paraformaldehyde 32% Solution, EM Grade | Electron Microscopy<br>Sciences | 15714-S |
| BLZ945 | Novartis | NA |
| BLZ945 (Sotuletinib) | MedchemExpress | HY-12768 |
| Eye spears | BVI Weck-Cel | 0008685 |
| Zinc Phosphate Cement Liquid | Hoffmann's | 8001300 |
| Zinc Phosphate Cement Powder | Hoffmann's | 80012-(* ) |
| Critical commercial assays |  |  |
| Deposited data |  |  |
| GEO |  |  |
| Experimental models: Cell lines |  |  |
| CT2A-Luc2-TdTomato |  |  |
| Experimental models: Organisms/strains |  |  |
| B6.Cg-Gt(ROSA)26Sortm9(CAG-tdTomato)Hze/J | Jackson stock no. | JAX: 007909 |
| B6.Cg-Gt(ROSA)26Sortm14(CAG-tdTomato)Hze/J | Jackson stock no. | JAX: 007914 |
| C57BL/6-Mrc1em1(cre/ERT2)Jkps/J | Jackson stock no. | JAX: 040118 |
| C57BL/6-Lyve1em1(cre/ERT2)Jkps/J | Jackson stock no. | JAX: 040119 |
| C57BL/6-Tg(Pf4-icre)Q3Rsko/J | Jackson stock no. | JAX: 008535 |
| B6.Cg-Csf1rtm1.2Jwp/J | Jackson stock no. | JAX: 021212 |
| C57BL/6-Tg(UBC-GFP)30Scha/J | Jackson stock no. | JAX: 004353 |
| B6.129P-Cx3cr1<tm1Litt>/J | Jackson stock no. | JAX: 005582 |
| B6.129S4-Ccr2tm1lfc/J | Jackson stock no. | JAX: 004999 |
| B6.129P2(Cg)-Cx3cr1tm2.1(cre/ERT2)Litt/WganJ | Jackson stock no. | JAX: 021160 |
| B6;129-Sall1<sup>tm5</sup>(cre/ERT2)Ryn | Prof. Nishinakamura<br>(Kumamoto University) |  |
| B6;129-Sall1<sup>tm4.1</sup>Ryn | Prof. Nishinakamura<br>(Kumamoto University) |  |
| B6.Cg-Spi1<sup>tm2</sup>Dgt/J | Jackson stock no. | JAX: 006922 |
| B6.129-Tg(Flt3-cre)#Ccb/leg | Timm Schroeder, D-BSSE Basel |  |
| B6.129S4-Ccr2<sup>tm1</sup>lfc/J | Jackson stock no. | JAX: 04999 |
| B6.Cg-Tg(S100A8-cre,-EGFP)1Ilw/J | Jackson stock no. | JAX: 021614 |
| CBA.Cg-Csf1<sup>rem1</sup>Prdns/J | Jackson stock no. | JAX: 032783 |
| Oligonucleotides |  |  |
| Software and algorithms |  |  |
| FlowJo v10.10 |  |  |
| QuPath | Bankhead et al. |  |
| R | R Core Team, 2024 |  |
| OMERO | Allan et al. |  |
| NEWTON v7 | Vilber, France |  |
| Other |  |  |
| Gammacell® 40 Exactor irradiator | Best Theratronics Ltd.,<br>Canada |  |

## EXPERIMENTAL MODEL AND STUDY PARTICIPANT DETAILS

### METHOD DETAILS

#### Animals

All animal handling, surveillance, and experimentation were performed according to the guidelines and legislation of the Swiss Federal Veterinary Office (SFVO) and the Cantonal Veterinary Office, Basel-Stadt, Switzerland, under license # 2929. Parabiosis experiments were performed at Washington University, St. Louis USA (J. Kipnis Lab). *Csf1r^fl/fl^* mice (B6.Cg-Csf1rtm1.2Jwp/J) were crossed to *Sall1^CreERT^*^2^ kindly provided by Prof. Nishinakamura (Kumamoto University) or to *Cx3cr1^CreERT^*^2^ mice, received from Prof. Niess (University of Basel) generating tamoxifen-inducible *Csf1r* knockout mice. TdTomato reporter mice (B6.Cg-Gt(ROSA)26Sortm9(CAG-tdTomato)Hze/J) were generated by crossing with the respective Cre-line, including *Sall1^CreERT^*^2^*, Cx3cr1^CreERT^*^2^*, Flt3^Cre^* (kindly provided by Timm Schroeder, D-BSSE Basel), *Pf4^Cre/wt^* (kindly provided by Prof. Skoda, University of Basel), *Mrc1^CreERT^*^2^ (kindly provided by J. Kipnis). *Sall1^GFP/wt^*were kindly received from Prof. Nishinakamura (Kumamoto University). Dual microglia and BAM reporter (*Sall1^GFP/wt^* and *Pf4^Cre/wt^;R26^TdTomato^*) were generated by crossing *Sall1^GFP/wt^* and *Pf4*^Cre/wt^;*R26*^TdTomato^ mice.

*Lyve1^CreERT2/wt^;R26^TdTomato^*(Ai9) were kindly provided by J. Kipnis (Washington University, St. Louis). *Cx3cr1^CreERT2/wt^;PU.1^fl/fl^;R26^TdTomato^*and *Cx3cr1^CreERT2/wt^;PU.1^wt/wt^;R26^TdTomato^*controls were kindly provided by Prof. Glass (LMU, München Germany). *CCR2^-/-^* mice were kindly provided by Dr. Cavelti-Weder (University of Basel).

#### Tamoxifen administration

Tamoxifen was dissolved in corn oil at a concentration of 20 mg/mL by shaking overnight at room temperature (RT) in light-protected tubes. Mice received tamoxifen at a dose of approximately 75 mg/kg body weight by intraperitoneal injection once every 24 h for five consecutive d. For adult mice, a standard injection volume of 100 µL tamoxifen solution per animal was used.

#### BLZ945 (CSF1Ri) Treatment

BLZ945 (Novartis) was dissolved in 20% Captisol prepared in PBS and freshly prepared immediately prior to treatment. Mice were treated by oral gavage at the indicated frequency in each experiment with BLZ945 at either 200 mg/kg body weight (BW) for the dihydrochloride (HCl) form or 169 mg/kg BW for the free base form, according to the manufacturer’s recommendation (Novartis). For a single depletion–repopulation cycle, mice received daily oral gavage of BLZ945 or vehicle control for 7 consecutive d, followed by a 7d repopulation phase unless otherwise indicated. For repetitive depletion–repopulation paradigms, each 7d depletion cycle was followed by either a 1-week or 3-week repopulation period, with mice sacrificed 4 weeks after completion of the final cycle.

For CSF1Ri in *Pf4^Cre/wt^:R26^TdTomato^* mice, BLZ945 (Sotuletinib, MedChemExpress) was used. BLZ945 was dissolved by repeated vortexing and sonication in a 37□°C water bath until fully resuspended. Mice were treated with BLZ945 at a dose of 200□mg/kg body weight.

#### Irradiation Experiment

All irradiations were performed using a Gammacell® 40 Exactor irradiator (Best Theratronics Ltd., Canada).

The head-protection chamber was designed (by Paul Jordi, Jordi Strahlentechnik, Münchenstein, Basel) with staggered lead wall offsets to reduce scatter and edge-associated radiation leakage. All chamber walls (side, roof, and bottom) consisted of 3□cm lead shielding. Because irradiation was delivered from both top and bottom sources, the roof and floor were reinforced with an additional 1.5□cm lead layer, resulting in a total shielding thickness of 4.5□cm in these regions.

Validation of the head-shielding efficiency was measured using individually packed thermoluminescent dosimeters (TLDs; detection range 0.1–10□Gy). TLDs were positioned on the exposed back and beneath the lead-protected head within the irradiation chamber to quantify radiation exposure under head-protected conditions relative to total body irradiation. An additional non-irradiated TLD was included as a baseline control. A total irradiation dose of 6.11□Gy was applied.

#### Bone marrow chimeras

Bone marrow chimeras were generated using a head-protected irradiation protocol. Recipient mice were anesthetized with ketamine/xylazine and placed into the irradiation chamber with the head shielded using a custom 3–4.5 cm lead barrier, as illustrated in Fig. S3B. Mice received two doses of 6 Gy total body irradiation separated by an interval of at least 4 h.

During the interval between irradiation cycles, bone marrow cells were isolated from donor mice under sterile conditions. Femurs and tibiae were collected and flushed with RPMI medium. Cell suspensions were filtered through a 70 µm filter and subjected to 1min ACK lysis. Cells were subsequently washed, resuspended in sterile PBS, and counted prior to transplantation. Recipient mice received 2 × 10^6^ sex-matched whole bone marrow cells by intravenous tail vein injection immediately following the second irradiation cycle.

For the assessment of clonal proliferation, we transplanted mixtures (1:1 ratio) of either *Cx3cr1*^CreERT2/wt^:*R26*^TdTomato^ and *Cx3cr1*^GFP/wt^ or *Mrc1*^CreERT2^:*R26*^TdTomato^ and *Cx3cr1*^GFP/wt^ BM into head-shielded *Cx3cr1*^CreERT2/wt^:*Csf1r*^fl/fl^ recipient mice.

#### Parabiosis

Parabiosis experiments were performed between *UBC^GFP^*mice and age- and sex-matched (female) *Sall1^CreERT2/wt^:Csf1r^fl/fl^ or Sall1^wt/wt^:Csf1r^fl/fl^* mice as previously described^54^. Briefly, mice were anesthetized with a ketamine/xylazine (100mg/kg, 10 mg/kg) mixture and the corresponding lateral sides were shaved and disinfected using alternating betadine and alcohol washes. A skin incision extending from the olecranon to the knee joint was generated, and subcutaneous fascia was carefully separated to create a free skin margin. Corresponding knee and elbow joints of parabiont pairs were connected using absorbable 5-0 Vicryl sutures, followed by approximation and closure of the dermis while excluding epidermal layers at the junction site. Animals received postoperative analgesia with buprenorphine (Buprenorphine SR; 1.0 mg/kg) and antibiotic supplementation in the drinking water.

#### Calvarium bone-flap transplantation

##### *Sall1*^Cret/wt^:*Csf1r*^fl/fl^

7–10-week-old mice were anesthetized with a ketamine/xylazine mixture (80 mg/kg ketamine, 16 mg/kg Xylazine) and maintained at 37°C on a heating pad throughout the surgical procedure. Hair over the head was removed using depilatory cream, and the surgical site was thoroughly disinfected with Betadine. A midline skin incision was made to expose the parietal and interparietal bones. A cranial window of approximately 4 × 6 mm was generated using an electric drill. During drilling, the surgical area was carefully monitored and kept moist as needed. The cranial flap was then gently removed by carefully detaching the underlying dura from the bone, while preserving the integrity of the dura mater, brain parenchyma, and sinus structures.

Following craniotomy, a sex- and age-mismatched fluorescent donor mouse, typically UBC-GFP, was euthanized. A donor calvarial flap of comparable size and from the corresponding skull region was prepared. The donor dura was removed from the bone flap, and the flap was trimmed as needed to match the recipient craniotomy site. The donor bone flap was then positioned over the cranial window and fixed in place using dental cement. Care was taken to prevent cement leakage between the fracture margins to allow proper bone apposition and healing. After the cement had hardened and the transplanted flap was stably immobilized, the skin was re-approximated and sutured.

Postoperative care included analgesic treatment without antibiotic administration. No postoperative infections were observed. Following surgery, mice were housed individually during the recovery period.

For GFP+ cell detection by HDFC, whole brains were processed and acquired. For immunofluorescence analysis, at least twenty 40-µm-thick sagittal brain sections within the transplantation area were analyzed per mouse.

##### Tumor cell injection for scRNAseq

7–15-week-old mice were anesthetized and maintained under anesthesia using isoflurane throughout the surgical procedure. Animals were mounted onto a stereotactic frame (Neurostar, Germany), and the scalp was disinfected using povidone-iodide solution prior to a midline incision. A burr hole was manually drilled 2□mm lateral to the cranial midline and 1□mm posterior to the bregma suture. Intracranial injections were performed using the StereoDrive v1 software (Neurostar, Germany) and a 10□µL Hamilton syringe (#80300, Hamilton, USA). The syringe was lowered to a depth of 3.0□mm below the dura surface and subsequently retracted by 0.5□mm to create a small reservoir. A total of 1.5 × 10^5^ CT2A-Luc2-TdTomato tumor cells suspended in 4□µL were injected at a rate of 1□µL/min. Following injection, the needle was left in place for at least 2□min and then slowly retracted in 50□µm incremental steps. The incision was subsequently sutured.

Tumor engraftment was confirmed on day 7 following intracranial implantation by bioluminescence imaging. Mice received an intraperitoneal injection of D-luciferin (150□mg/kg^-1^), and imaging was performed 10□min after injection using the NEWTON 7.0 instrument (Vilber, France) and bioluminescence counts measured using NEWTON v7 Software (Vilber, France) imaging system and software (Vilber, France).

#### Mouse tissue collection and processing

##### CNS tissue

For all mouse cohorts, animals were deeply anesthetized by intraperitoneal injection of ketamine/xylazine (80□mg/kg ketamine [Ketanarkon, Streuli Tiergesundheit, Switzerland] and 16□mg/kg xylazine [Rompun, Elanco, USA]) and transcardially perfused with PBS prior to tissue collection. Following perfusion, the dura mater was carefully peeled from the cranium, the fourth ventricle was removed, and brains were immediately processed for downstream applications. Brain hemispheres were separated, with one hemisphere immersion-fixed in 4% paraformaldehyde (PFA) for histological and immunofluorescence analyses, while the contralateral hemisphere was used for flow cytometric analyses following dissection of the lateral ventricle choroid plexus. Brain tissue was subsequently minced and maintained in ice-cold PBS until processing.

All tissues were enzymatically digested at 37□°C for 30□min using 1□mg/mL collagenase type IV (#LS004188, Worthington Biochemical Corporation, USA) and 250□U/mL DNase I (#10104159001, Roche, Switzerland) in digestion buffer consisting of HBSS with Ca^2+^/Mg^2+^ (#14205-050, Gibco, USA), 1% MEM non-essential amino acids (NEAA), 1□mM sodium pyruvate, 44□mM sodium bicarbonate (#25080-060, Gibco, USA), 25□mM HEPES (#H0887, Gibco, USA), 1% GlutaMAX-I, and 1% antibiotic-antimycotic (#15240062, Gibco, USA). Following digestion, tissues were mechanically dissociated by repeated trituration using a 1000□µL pipette tip. Brain-derived cell suspensions were subsequently filtered through a 70□μm cell strainer and subjected to myelin and debris removal using debris removal solution (#130-109-398, Miltenyi Biotec, Germany) according to the manufacturer’s instructions. Cells were washed with PBS and maintained on ice until further downstream applications.

Dura and choroid plexus samples were processed using the identical enzymatic digestion protocol; however, no myelin/debris removal step was performed. Following enzymatic dissociation, cell suspensions were filtered through a 35□μm mesh cell strainer cap (Falcon®, #352235) prior to downstream flow cytometric analyses.

##### Blood

Mice were gently restrained and tail disinfected with 70% ethanol. Peripheral blood was collected from the lateral tail vein by puncture and immediate collection of 2 × 10□µL blood into a 96-well plate containing 100□µL PBS supplemented with 1□mM EDTA and 1:100 mouse FcBlock (TruStain FcX (#101320, BioLegend, USA)).

#### Spectral flow cytometry and FACS

##### Staining and acquisition

All samples were acquired on a Cytek Aurora 5-Laser Spectral Analyzer (Cytek Biosciences, USA) using standard daily quality-controlled Cytek-Assay-Settings unless otherwise stated.

##### CNS tissues

Single cell suspensions were resuspended at equal volumes in PBS and distributed in 96-well plate (round bottom). All centrifugation and incubation steps were performed at 300□×□g at 4□°C, protected from light, unless otherwise stated. Viability staining was performed by incubating cells with a Zombie Viability dye (Zombie Aqua or NIR Viability kit) for 15□min. Subsequently, Fc-block was performed by incubating cells in a 1:50 dilution of mouse TruStain FcX (#101320, BioLegend, USA) for 15□min. Antibody mastermixes (full-stains and FMOs) were freshly prepared on the day of staining in Brilliant Stain Buffer (#00-4409-75, ThermoFisher Scientific, USA). Cell surface markers were stained with surface antibody mastermixes for 20□min and then washed three times in autoMACS Running Buffer. Fixation/permeabilization step was performed by incubating the cells for 20□min at RT using a Cyto-Fast Fix/Perm Buffer set (#426803, BioLegend, USA). Finally, the samples were washed twice, resuspended in a final volume of 200□µL of autoMACS Running Buffer, and acquired. Absolute counts were quantified by normalizing cell count by acquisition volume.

##### Blood

All centrifugation and incubation steps were performed at 400□×□g at 4□°C, protected from light, unless otherwise stated. For longitudinal studies of blood over multiple days, reagents (antibody mastermixes, FMO mastermixes, viability dyes and FcBlock-EDTA mix) was prepared in advance. To correct for potential batch effects, blood of a C57BL/6 mouse was collected transcardially into 1mM EDTA supplemented with 1:100 FcBlock and frozen at −80 °C prior to experiment start. For each timepoint, 1 vial of blood was thawed and stained side by side with other samples and used to assess whether batch normalization is required. After blood collection as described in “Mouse tissue collection and processing”, blood was centrifuged and resuspended in Zombie Viability dye (Zombie Aqua or NIR Viability kit) for 15 min. Subsequently, cell surface markers were stained with surface antibody mastermixes for 20□min and washed three times in autoMACS Running Buffer. Then, blood was fixed/lysed by adding 150 µL of BD FACS Lysing Solution (BD Biosciences, #349202) for 10 min at RT. Cells were centrifuged at 600□×□g at 4□°C, washed twice and resuspended in 200 µL autoMACS running buffer (20 µL initial whole blood resuspended in 200 µL acquisition volume) and subsequently acquired on a Cytek Aurora 5-Laser Spectral Analyzer (Cytek Biosciences, USA). Absolute counts were quantified by normalizing cell count by acquisition volume.

##### Flow chytometry analysis

For conventional flow cytometry analysis, frequencies/counts, acquisition volume/sample and median fluorescence intensities (MFI) of gated populations were exported along corresponding metadata and analyzed in R.

##### FlowSOM analysis

Data was unmixed using the SpectroFlo software (Cytek) and then manually pre-gated on CD45^+^/CD11b^+^, live, single cells and then cells of interest (i.e CD45^+^CD11b^+^Ly6C^-^CD64^+^ macrophages) using FlowJo v10 Software. The analysis was subsequently performed in R v4.3.2. Data was transformed by arc-sinh transformation using variance stabilizing cofactors for each channel (estParamFlowVS and transFlowVS functions from the FlowVS package). Manual corrections of cofactors were applied when necessary. Clustering was performed using FlowSOM and ConsensusClusterPlus using the wrapper function cluster from the CATALYST package. The resulting clusters were manually annotated and merged based on marker expression. Clustering identity was compared to conventional gating by projection of cluster identity onto bivariate plots. For visualization and statistical quantification, single cell object was transformed into a dataframe and analysed as indicated in respective figure legends.

#### Fluorescence-activated cell sorting

##### FLASHseq of TdTomato+ macrophages

Samples and collection plates were maintained at 4□°C throughout the tissue preparation sorting procedure, and PBS was used as the sorting and sample preparation buffer in all steps. Tissue was prepared as described in “Mouse tissue collection and processing”. Viability staining was performed using Zombie NIR at 1:1000 dilution in PBS for 20 min. No FcBlock was perfomed as no surface staining was required.

Pf4-TdTomato^+^ live singlet cells from brain and dura preparations were single-cell sorted on a BD FACSDiscover S8 Cell Sorter using a 100□µm nozzle into lysis buffer-prefilled Eppendorf twin.tec® 384-well PCR plates (Cat. No. 0030128508, Eppendorf) mounted on a cooled plate holder, with one cell deposited per well. Plate was immediately sealed and centrifuged at 700 □×□g at 4□°C and transferred into −80□°C. scRNA-sequencing libraries were generated according to the FLASH-seq protocol (v4).

##### 10X scRNAseq of tumor engrafted mice

Samples and collection tubes were maintained at 4□°C throughout the tissue preparation, sorting procedure, and PBS was used as the sorting and sample preparation buffer in all steps Mice were deeply anesthetized and perfused as described in “Mouse tissue collection and processing”. Brain was split into tumor-bearing and tumor-free hemisphere by mid-sagittal cut and processed as described in “Mouse tissue collection and processing”. ACK lysis was performed once for 5 min. Cells were Fc-blocked and stained with anti-CD45 and anti-CD11b for 30 min at 4□°C. Cells were washed and Hashtag-antibodies added followed by 20 min incubation. Samples were washed and then pooled before the final wash and subsequent resuspension in PBS and 0.02% BSA. DAPI was added for dead cell exclusion.

#### Immunofluorescence

##### Mouse samples

Mice were transcardially perfused with PBS. Brains were subsequently dissected and post-fixed overnight in 4% paraformaldehyde (PFA) at 4 °C, followed by thorough washing in PBS. Dura mater and choroid plexus samples were fixed in 4% PFA for 10 min at room temperature prior to washing in PBS. Brain tissue was sectioned using a vibratome (40 µm sagittal sections and 27 µm coronal sections) and processed as free-floating sections. Choroid plexus and isolated pia preparations were similarly stained free-floating. Dura samples were carefully dissected under a stereomicroscope and stained on glass slides. All samples were blocked and permeabilized in PBS containing 3% bovine serum albumin (BSA) and 0.5% Triton X-100, followed by incubation with primary antibodies overnight at 4 °C. After washing, appropriate fluorescent secondary antibodies were applied for 1 h at room temperature, followed by additional washing steps prior to mounting.

#### Human samples

Human tissue samples obtained from healthy and Alzheimer donors were formalin-fixed and paraffin-embedded (FFPE). Tissue blocks were sectioned at 5 µm using a Microm HM 355S microtome (Thermo scientific) and mounted onto glass slides (Epredia). Immunofluorescence was performed with the Ventana Discovery Ultra (Roche Diagnostics Suisse SA) automated stainer. In brief, tissue sections were deparaffinized and rehydrated. Antigens were retrieved by heat in Cell Conditioning buffer 1 (CC1, ref. 950-124, Ventana) at 98°C for 40 minutes. Primary antibody against IBA1 (goat monoclonal, 1:500; Novus Biologicals) and CD163 (rabbit monoclonal, 1:150; Abcam) was manually applied and incubated for 1 hour at 37°C. After washing, secondary antibodies donkey anti-goat Cy3 (Jackson ImmunoResearch) and donkey anti-rabbit Alexa Fluor 647 (Jackson ImmunoResearch) were applied and incubated for 1 hour at 37°C. Nuclear counterstaining was performed using DAPI, incubated at RT for 30 min. All the secondary antibodies were used at a concentration of 1:1000. After staining, slides were washed three times with 1×PBS and mounted using ProLong™ Gold Antifade Mountant (Thermo Fisher Scientific).

#### Microscopy

##### Stained slides

Fluorescence imaging was performed using inverted spinning-disk confocal microscopy systems (Nikon Eclipse Ti and Nikon Ti2; Nikon). High-resolution imaging was acquired using a 40× air objective, while lower magnifications (4×, 10×, and 20×) were used for overview and large-area imaging.

Images were acquired using standard filter sets appropriate for the respective fluorophores. Z-stacks and tiled acquisitions were performed where indicated.

For quantitative analyses, imaging settings (including laser power, exposure time, and detector gain) were kept constant within each experiment for samples stained and imaged under identical conditions.

##### Whole leptomeninges microscopy

Whole-pia imaging was performed on intact brains using spinning-disk confocal microscopy (Nikon Ti2) with a 4× objective. Z-stack tile scans were acquired across the entire pial surface of the dorsal cortex in ice-cold PBS prior to downstream processing.

Due to the autofluorescence of the leptomeninges, signal detection was effectively restricted to the surface, with minimal contribution from underlying cortical tissue, as verified by three-dimensional reconstructions.

Equivalent acquisitions were performed on fixed hemispheres or whole brains for structural comparison.

##### Tissue clearing and lightsheet microscopy

Brains were cleared using EZ Clear with refractive index matching prior to imaging. Cleared brain hemispheres were immersed in oil and imaged using a benchtop mesoSPIM system equipped with a motorized XY stage and controlled via mesoSPIM-control software. Imaging was performed using a 5×/0.14 NA air objective (working distance 3.4 mm) and a Photometrics IRIS 14 camera.

tdTomato fluorescence was excited at 561 nm and detected using a 595/50 bandpass emission filter. Dual-sided illumination was used to reduce shadowing artifacts and improve signal uniformity.

Tile scans and z-stacks were acquired to cover entire hemispheres. Image stitching was performed in Fiji, and three-dimensional visualization was carried out using Imaris.

#### ScRNAseq data analysis

##### BAM-Atlas

Raw count matrices for four wild-type samples from Van Hove et al. (GSM3687213–GSM3687216; dura mater, choroid plexus, enriched subduralmeninges, and whole brain) were obtained from NCBI GEO. Cells were filtered by MAD-based outlier detection (nmads = 4) on detected gene count, total UMI count, and mitochondrial read fraction, followed by PCA based outlier removal on QC metrics (scater). Genes with average count < 10□³ were excluded. Non-myeloid populations (dendritic cell subsets, B cells, T cells, NK cells, ILC, neutrophils, and doublets) identified at Louvain resolution 0.6 were removed, retaining microglia, macrophages, and monocytes.

The myeloid subset was normalised using SCTransform, regressing out dissociation stress score, cell cycle scores (S and G2/M phase), and ribosomal RNA fraction. Datasets were integrated across tissues using Seurat canonical correlation analysis (CCA; SelectIntegrationFeatures, nfeatures = 3,000; PrepSCTIntegration; FindIntegrationAnchors; IntegrateData). PCA was run on the integrated assay; UMAP (dims = 1:15, n.neighbors = 20) and tSNE (dims = 1:15) were computed on the integrated space. Louvain clustering was performed at resolutions 0.1–1.2; resolution 0.8 was used for annotation. Clusters were manually assigned to cell types based on canonical marker expression, and contaminating populations (B cells, mast cells, endothelial, stromal, CD45□, and unassigned cells) were removed to produce the final annotated object. Differentially expressed genes between BAM subpopulations were identified using FindMarkers (Seurat, SCT assay, min.pct = 0.25, logfc.threshold = 0.4).

##### Tumor-bearing and tumor-free hemisphere 10X scRNAseq

CD45□ cells were sorted from CT-2A glioma-bearing mouse brains across four sample groups (Csf1r-Cre□□ × tumour contralateral hemisphere) and sequenced on a NovaSeq SP lane (800M reads). Reads were aligned to mm10Ensembl 102 using STARsolo. Cells were filtered, 600 UMIs and 450 detected genes; doublets were removed in two steps using HTO classification and scDblFinder. Batch effects across sample groups were corrected using FastMNN (batchelor; k = 20, d = 50). Louvain clustering was performed on the MNN-corrected space at k = 10; all visualisations use UMAP.MNN (n_neighbors = 15) and TSNE.MNN embeddings. Cell-type annotation combined SingleR (ImmGenData; Philip Schmassmann dataset ^70^) with manual marker-based assignment. Cluster 1 (MG-like cells) was re-clustered independently; subclusters were characterised using scoreMarkers (scran), and gene sets defined by markers with mean AUC >0.65 were projected onto the M-Verse multi-dataset integration using AddModuleScore (Seurat). For Figure 6D, pseudobulk differential expression was computed in MG-like and Microglia_1 clusters using limma on aggregated counts (≥ 20 cells per pseudobulk), with the contrast tumor-free hemisphere Cre+ versus tumor-free hemisphere Cre-.

##### FLASHseq

Cells sorted from the brain and dura mater of Pf4^Cre/wt^;R26^tdTomato^ wild-type mice, and from the brains of Cx3cr1^GFP/wt^ and Cx3cr1CreERT2/wt;R26^tdTomato^ mice, as described above, were processed using the FLASHseq protocol (768 single-cell libraries). FLASH-seq libraries were sequenced on an Illumina NextSeq 500/550 platform to generate 36-bp single-end reads. A custom reference genome was assembled from mm39 supplemented with eGFP, eYFP, CreERT2, and TdTomato transgene sequences and indexed with Ensembl 110 annotation. Reads were aligned using STAR and gene-level counts were quantified with featureCounts within Snakemake workflows, executed on the SciCore HPC cluster. The resulting count matrix was imported into Seurat (min.cells = 3, min.features = 300) and filtered (detected features > 300 and < 7,500, UMI count > 300, mitochondrial fraction < 8%). Data were log-normalised (scale factor = 10,000), highly variable features identified (vst, nfeatures = 2,000), and UMAP computed on the first 25 PCs. Cells were classified as BAMs (UMAP_2 > 8) or microglia (UMAP_2 ≤ 8) based on UMAP coordinate position. Differential marker genes between BAMs and microglia were identified using scoreMarkers (scran), performed on the Pf4-Cre × Ai14 wild-type subset.

##### 3xDR

Raw count matrices for five samples (GSM7070553–GSM7070557; three control, two 3xDR) were obtained from NCBI GEO (GSE226286). Cells were filtered (detected features ≥ 200 and ≤ 4,000; mitochondrial fraction ≤ 5%) and doublets removed using scDblFinder (per-sample, with cluster-guided detection). After doublet exclusion, dimensionality reduction was recomputed on singlets (11 PCs, n.neighbors = 26). Macrophage clusters were identified at Louvain resolution 0.6, extracted, and independently re-clustered (12 PCs, n.neighbors = 9). Final cell-type annotations were assigned at resolution 0.4. Differential expression was performed using a pseudobulk strategy: counts were aggregated per mouse (make_pseudobulk()), and edgeR quasi-likelihood F-tests (filterByExpr, calcNormFactors, estimateDisp, glmQLFit, glmQLFTest) were applied. Significant genes were defined at FDR < 0.05 and |log□FC\| > 0.5. The primary comparisons shown in the figures were: BAMs 3xDR versus BAMs Control (Figure 5 volcano), and each repopulating subtype individually versus MG Control (IFN repop. Mac, MHCII^high^ repop. Mac, DAM^like^ repop. Mac; Venn, top 200 upregulated genes per subtype). To assess BAM identity re-expression, BAMs (Control) and repop_BAMs (3xDR) were each compared against the same background comprising all remaining non-BAM cell populations using Wilcoxon rank-sum tests (FindMarkers, logfc.threshold = 0, min.pct = 0.1); genes were classified as shared upregulated, shared downregulated, BAM-specific, or repop_BAM-specific based on adjusted p-value < 0.05 and log□FC directionality, and restricted to genes detected in > 25% of repop_BAM cells. Module scores for MHCII-high, DAM, and homeostatic MG signatures were computed using AddModuleScore (Seurat) and visualised as feature plots and dotplots.

###### ROSMAP

Single-nucleus RNA-sequencing data from the Religious Orders Study and Memory and Aging Project (ROSMAP) multiregion cohort were obtained from the Synapse data repository (syn52383412). Gene expression matrices were downloaded for six brain regions: angular gyrus (AG; syn52408594), entorhinal cortex (EC; syn52408588), hippocampus (HC; syn52408592), midtemporal cortex (MT; syn52408595), prefrontal cortex (PFC; syn52408599), and thalamus (TH; syn52408586). Cell type annotations were derived from the original study (Mathys, Boix et al., Nature 2024). From each regional dataset, nuclei annotated as CNS-associated macrophages (CAMs) or microglia (Mic; including P2RY12+, MKI67+, DUSP1+, and TPT1+ subtypes) were extracted using Seurat v5, yielding a combined dataset of 104,181 nuclei (CAMs: 4,194; Mic: 99,987) across all six regions.

The six regional subsets were merged into a single Seurat object with region-specific cell barcode prefixes. Highly variable features were identified using FindVariableFeatures (nfeatures = 2,000), followed by data scaling and principal component analysis (PCA) on the variable feature set. UMAP dimensionality reduction and shared nearest-neighbour (SNN) graph construction were performed on the top 20 principal components with 15 neighbours (RunUMAP: dims = 1:20, n.neighbors = 15; FindNeighbors: dims = 1:20). Unsupervised Louvain clustering was applied across a resolution range of 0.1–0.9 in steps of 0.1, and cluster stability was evaluated using clustree v0.5. Resolution 0.6 was selected as it provided biologically coherent clusters without over-fragmentation. Neuropathological Alzheimer’s disease status was derived from the pathAD metadata field, dichotomised as AD or non-AD.

To quantify enrichment of our murine peripheral macrophage gene signature (n = 96 genes; intersection of three independent murine repopulating-macrophage gene sets; see analysis in 3xDR above, converted to human orthologues) across clusters, we applied fast gene set enrichment analysis (fgsea v1.32.4). The gene signature comprised the intersection of three independent peripheral macrophage repopulation gene sets (top 200 genes each), retained only if present in the expression matrix. For each cluster, all expressed genes were ranked by their mean expression z-score (computed gene-wise from AverageExpression across clusters), such that genes most specifically expressed in a given cluster rank highest. fgsea was run with scoreType = “pos” to test exclusively for positive enrichment with nPermSimple = 10,000 permutations and a random seed of 42 for reproducibility. Normalised Enrichment Scores (NES) and nominal p-values were computed per cluster; multiple testing correction was applied globally across all cluster tests using the Benjamini-Hochberg (BH) procedure. Clusters with adjusted p-value < 0.05 were considered significantly enriched. Results are presented as a lollipop plot (NES on x-axis, point size proportional to −log10(padj), line style indicating significance).

To identify human BAMs, we curated a BAM gene signature from the study by^56^. Specifically, marker genes reported for BAM clusters 19 and 26 in Supplementary Table 3 were combined into a single non-redundant gene set. This curated signature was used to compute per-cell module scores using Seurat’s AddModuleScore function.

To identify transcriptional features specific to candidate clusters, pairwise differential expression was performed between each candidate cluster and a MG background using the Wilcoxon rank-sum test (FindMarkers, Seurat v5; min.pct = 0.10, log₂FC threshold = 0.25, Benjamini-Hochberg correction). C6 was compared against all clusters excluding C10 and C12, and C12 against all clusters excluding C6 and C10. Results are displayed as volcano plots with genes significant at p_adj < 0.05 and |log₂FC| ≥ 0.5. Top 10 up- and downregulated genes and a custom set of differentially expressed genes is displayed.

For clusters C6 and C12, the proportion of nuclei contributed by each brain region was calculated separately in AD and non-AD donors, with all cells in the cluster per diagnostic group as the denominator, and displayed as stacked proportional bar charts.

All analyses were performed in R (version 4.5.1). Key packages used include: Seurat (≥ 5.0), SingleCellExperiment, scran, scater, scDblFinder, edgeR, limma, batchelor (FastMNN), ComplexHeatmap, and the tidyverse suite. Snakemake pipelines were executed on the SciCore HPC cluster at the University of Basel

#### Histology

##### Quantification of cells

Cell detection and quantification were performed in QuPath using custom Groovy scripts. Cells were identified using the built-in watershed-based detection algorithm on the relevant marker channel (e.g., Iba1). Detected cells were classified into marker-defined populations based on fluorescence intensity thresholds in additional channels. Classification thresholds were defined empirically and kept constant within each dataset. Detection parameters were adjusted where necessary to account for differences in cell morphology between depleted and non-depleted conditions. For selected analyses, fluorescence intensity and morphological features were extracted.

##### Separation of leptomeninges and parenchymal/perivascular

Leptomeningeal and parenchymal/perivascular cells were distinguished based on spatial proximity to the tissue boundary. The tissue surface was defined from autofluorescence signal, and the distance of each detected cell to this boundary was calculated. Cells located within 5 µm of the surface were classified as leptomeningeal, while all others were assigned to parenchymal/perivascular compartments.

#### Monte Carlo spatial analysis

Spatial organisation of labelled cells was analysed in R using a Monte Carlo label-permutation approach based on QuPath-derived centroid coordinates. Analyses were performed separately for leptomeninges, dura, choroid plexus, and brain regions. For each region, neighbour densities for same-class cell pairs were calculated in concentric annuli up to 250 µm using 5 µm radial bins.

To generate the null distribution, cell-class labels were randomly permuted within each region while keeping cell coordinates fixed, thereby preserving tissue geometry and overall cell density. This was repeated 1,000 times. Observed neighbour-density curves were then compared with the permutation-derived null distribution using global simulation envelopes. Spatial deviations between observed and randomised distributions were additionally summarized over the 20–80 µm range.

#### Dot maps

Spatial distributions of labelled cells were visualized as dot maps based on centroid coordinates extracted from QuPath. Cells were plotted by class for individual regions, with coordinates displayed in a two-dimensional plane and axes scaled equally. Representative regions or all regions were visualized depending on the analysis.

#### Cell lines

The murine malignant astrocytoma cell line CT-2A was kindly provided by Prof. Thomas Seyfried (Boston College). CT-2A cells were cultured as adherent monolayers in high-glucose DMEM without glutamine (Gibco) supplemented with 10% fetal bovine serum (PAN-Biotech), 1% GlutaMAX (Gibco), and 1% penicillin–streptomycin (Sigma-Aldrich). Cells were maintained at 37□°C and 5% CO_2_ and passaged at ∼80% confluency. All cell lines were routinely tested and confirmed negative for mycoplasma contamination using a MycoAlert detection kit (#LT07-318, Lonza). CT-2A Luc2-tdTomato was produced by lentiviral transduction to express luciferase 2 (Luc2) and the tdTomato reporter protein as previously reported^70^.

#### Mouse genotyping

Genotyping primer list can be found in the table “Primer List” in the supplementary information.

### QUANTIFICATION AND STATISTICAL ANALYSIS

Data are presented as mean ± standard error of the mean (SEM) unless otherwise indicated. The exact number of biological replicates (n = mice), number of independent experiments and statistical tests used are specified in the corresponding figure legends. Statistical significance was defined as p < 0.05 (*p < 0.05, p < 0.01, ***p < 0.001, ****p < 0.0001; ns, not significant). All statistical analyses were performed using R software (v4.3.2). Given the typical sample sizes used in experimental biology studies, formal assessment of normality was not systematically performed. Parametric or rank-based non-parametric statistical tests were selected as appropriate for the experimental design and data distribution. Comparisons between two groups were performed using two-tailed unpaired or paired t tests, or rank-based non-parametric tests where appropriate. Multiple testing correction was applied where appropriate. All graphical illustrations were created with BioRender.com

