## Supplementary figures and images for "The brain-meningeal interface functions as a reservoir and entry site for brain parenchymal macrophages"

### Supplementary Figure 1

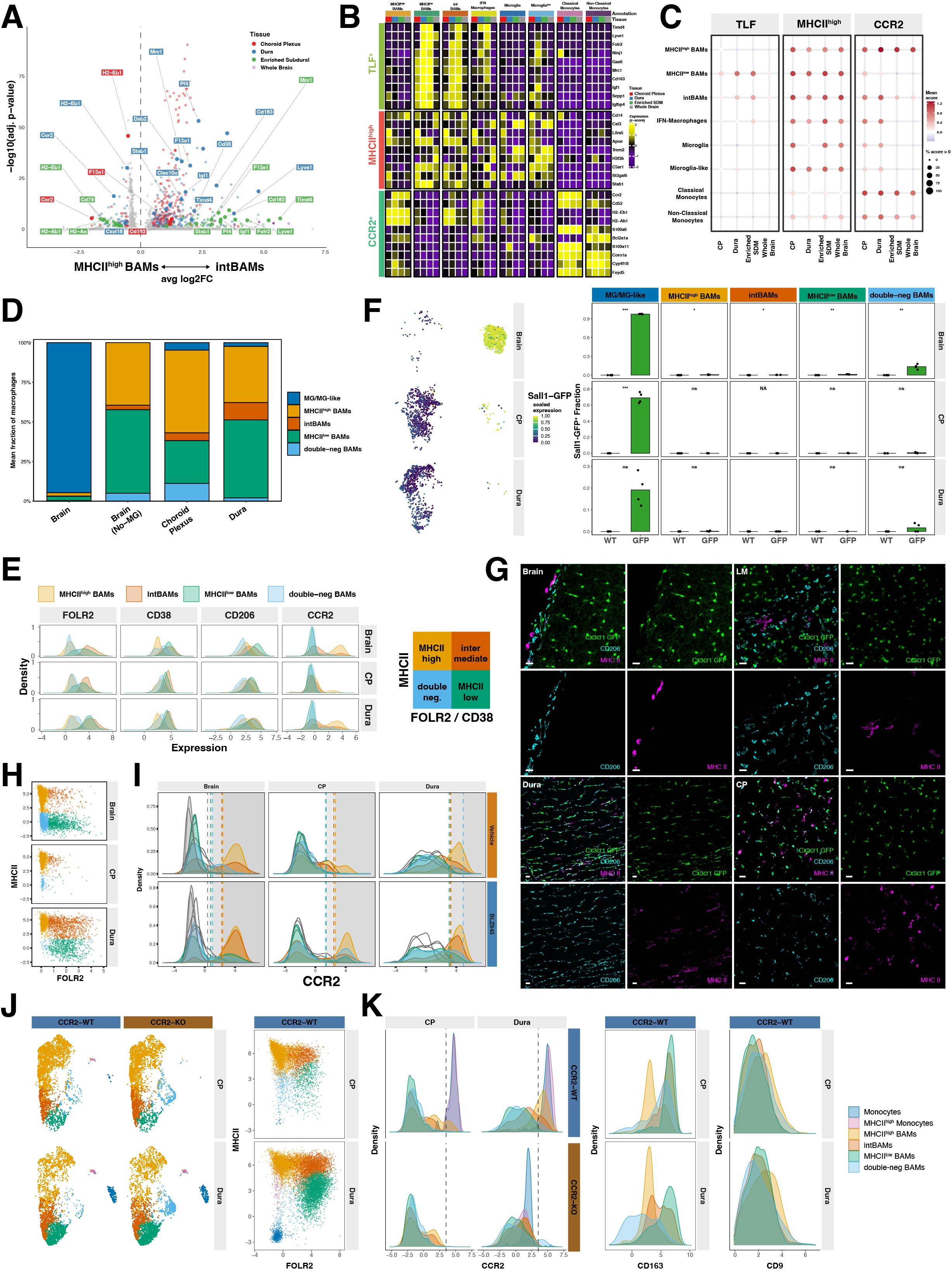

### Supplementary Figure 2

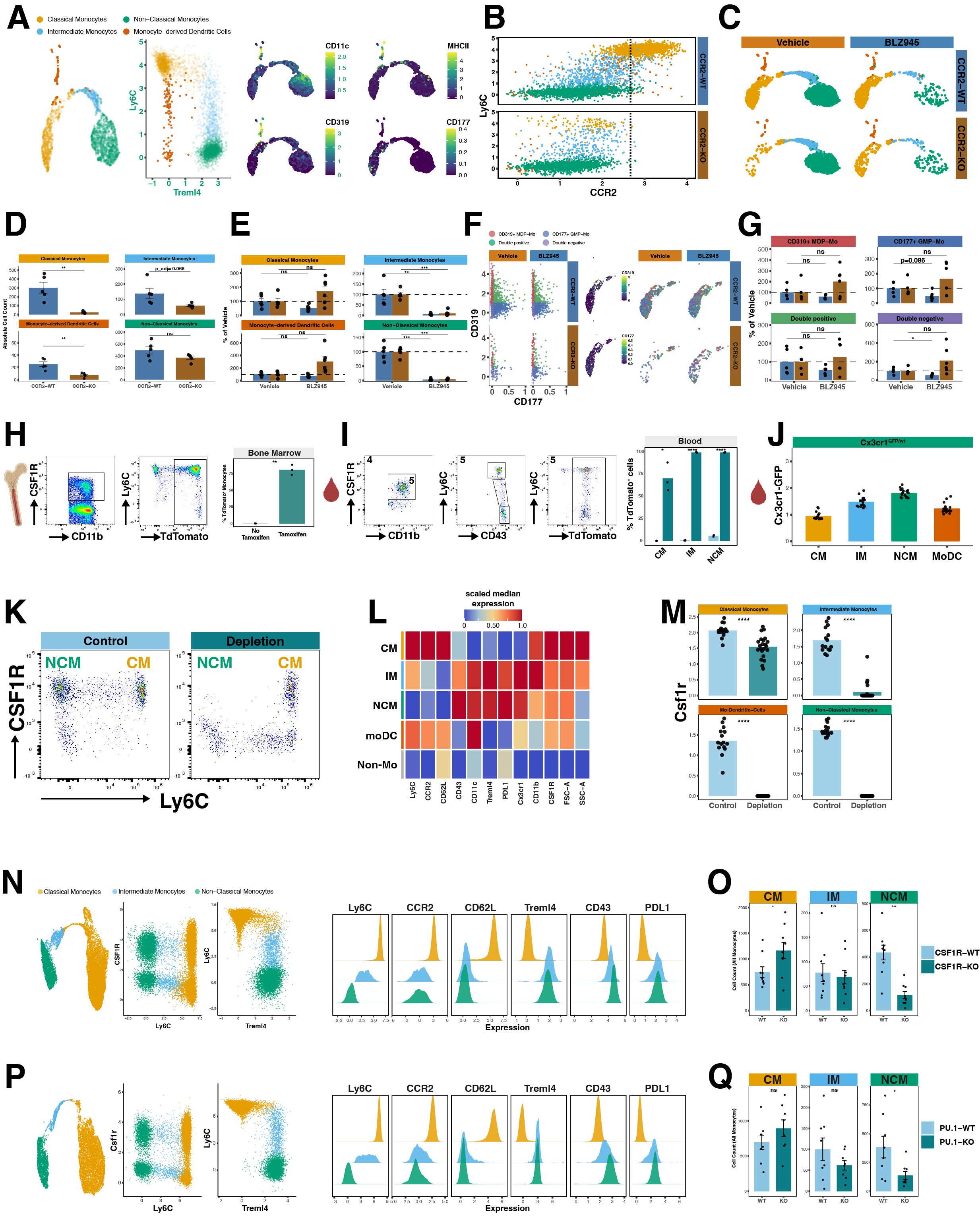

### Supplementary Figure 4

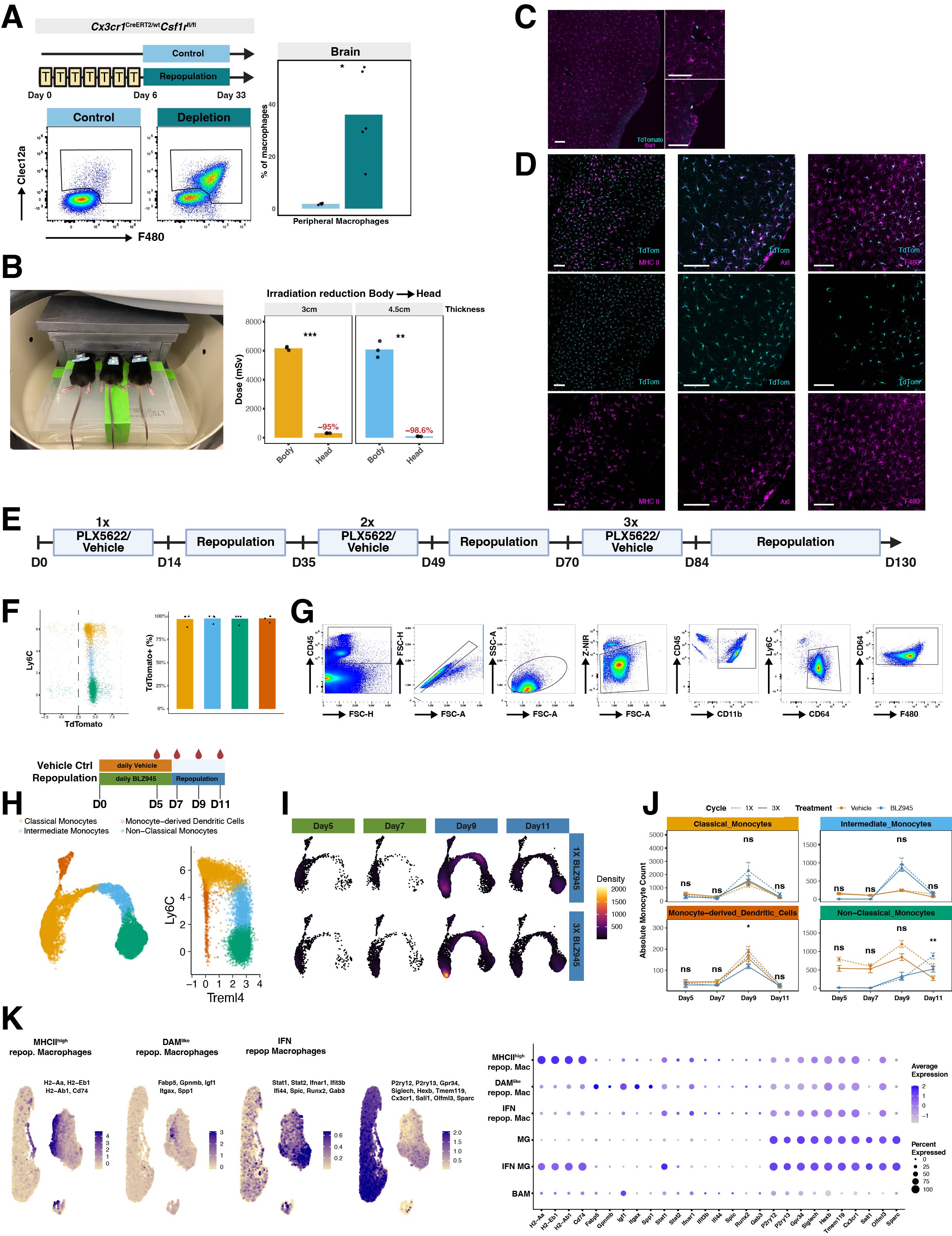

### Supplementary Figure 4

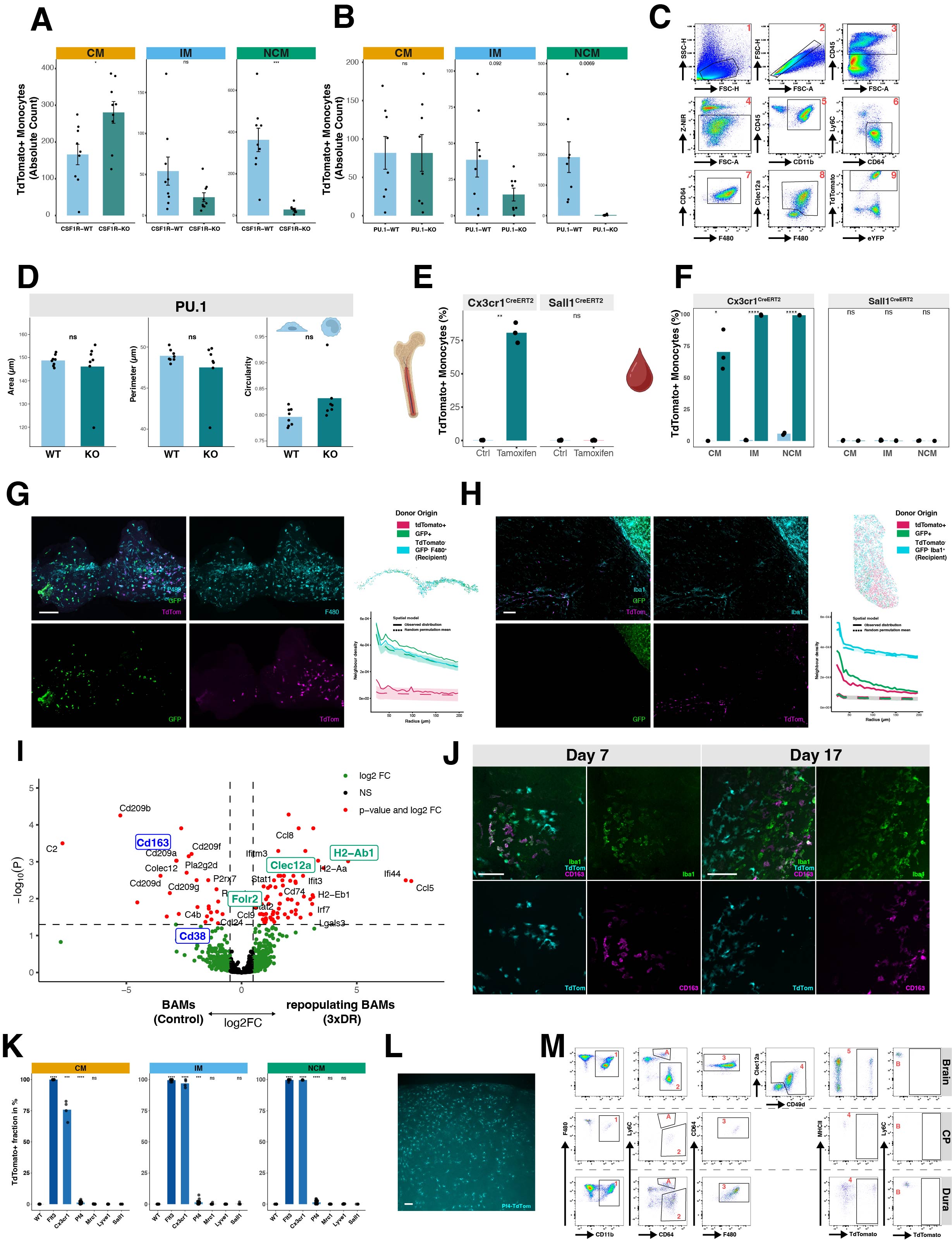

### Supplementary Figure 5

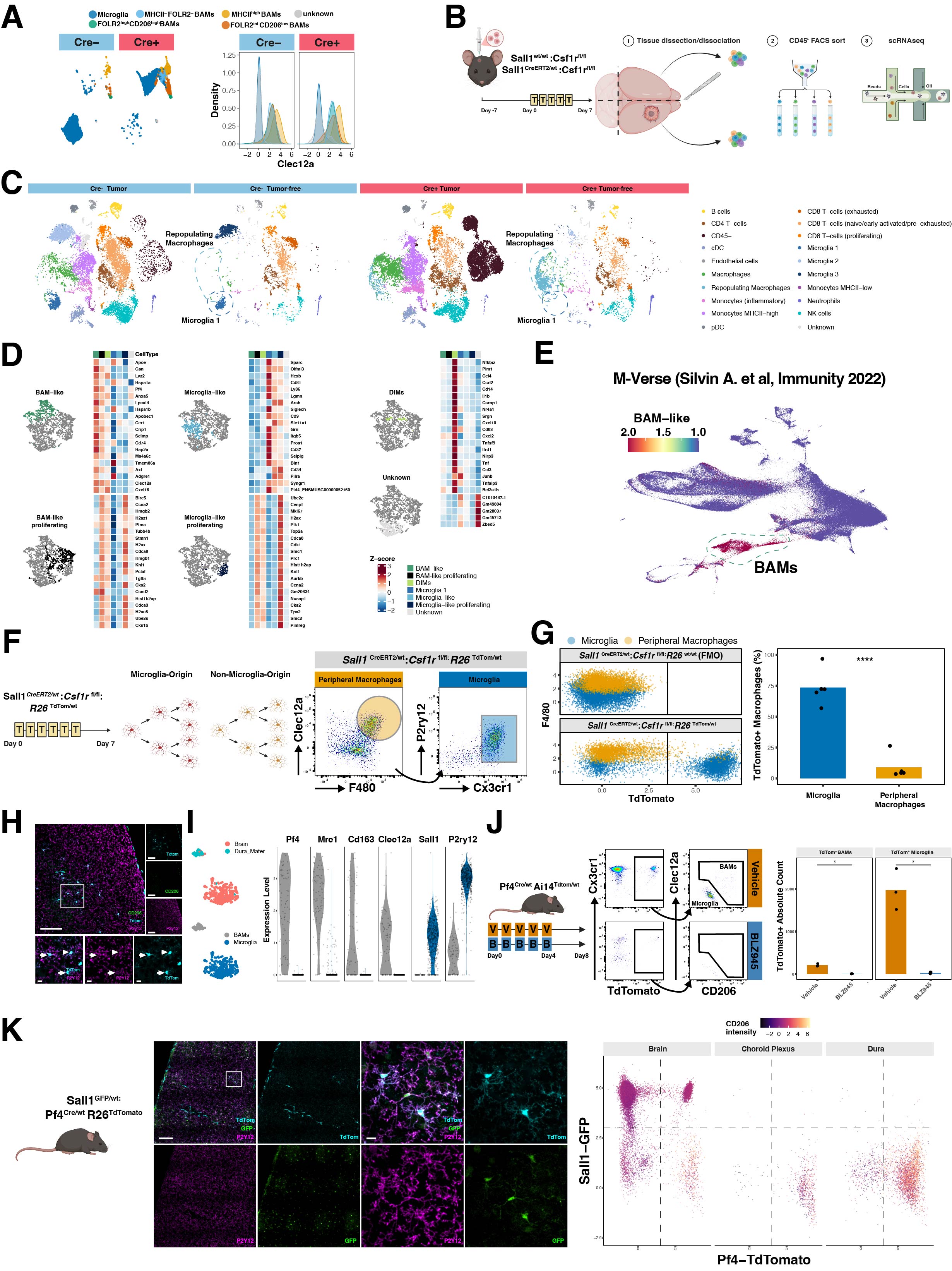

### Supplementary Figure 6

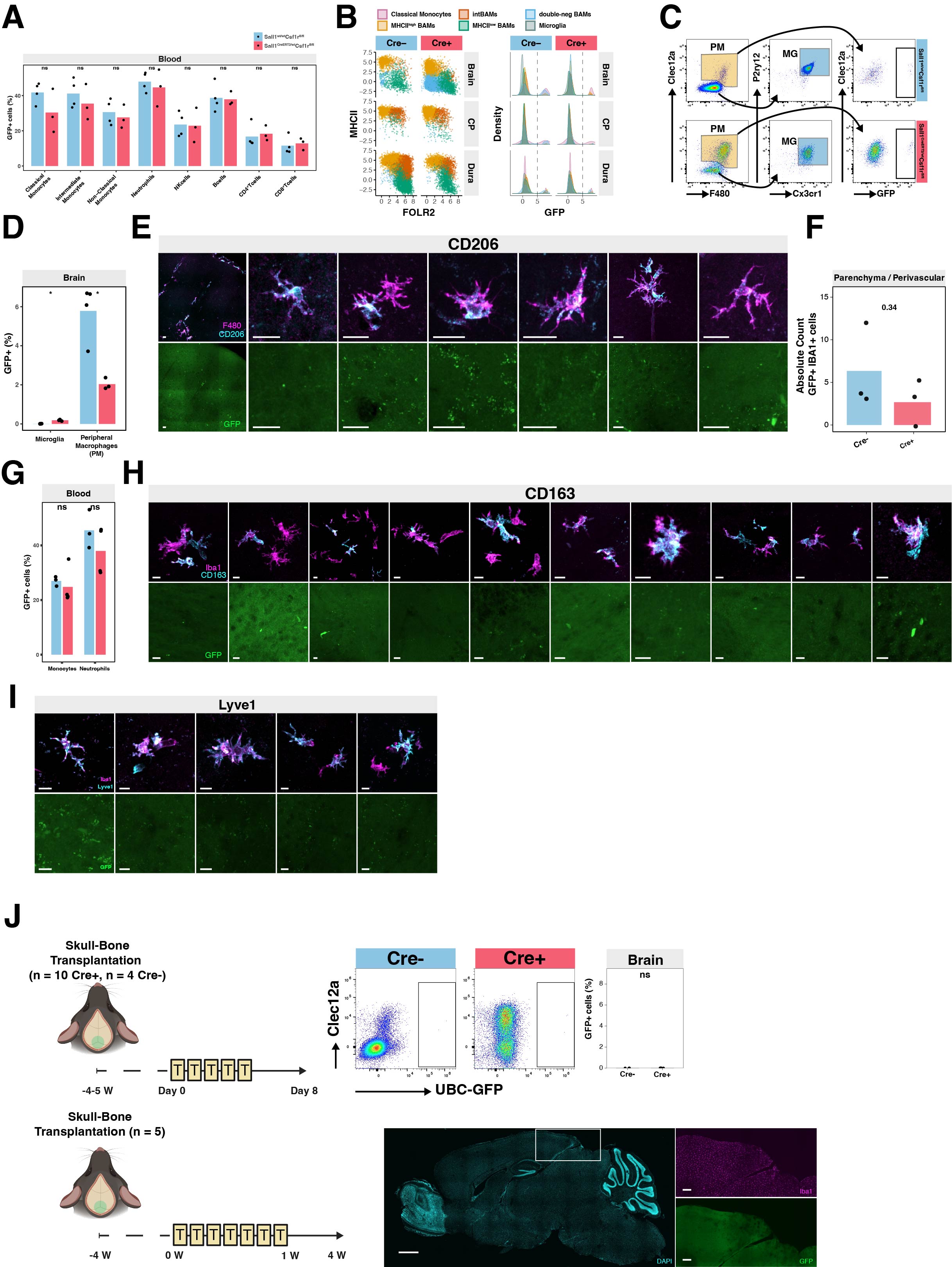
