## Supplementary material for "The brain-meningeal interface functions as a reservoir and entry site for brain parenchymal macrophages": Graphical Abstract

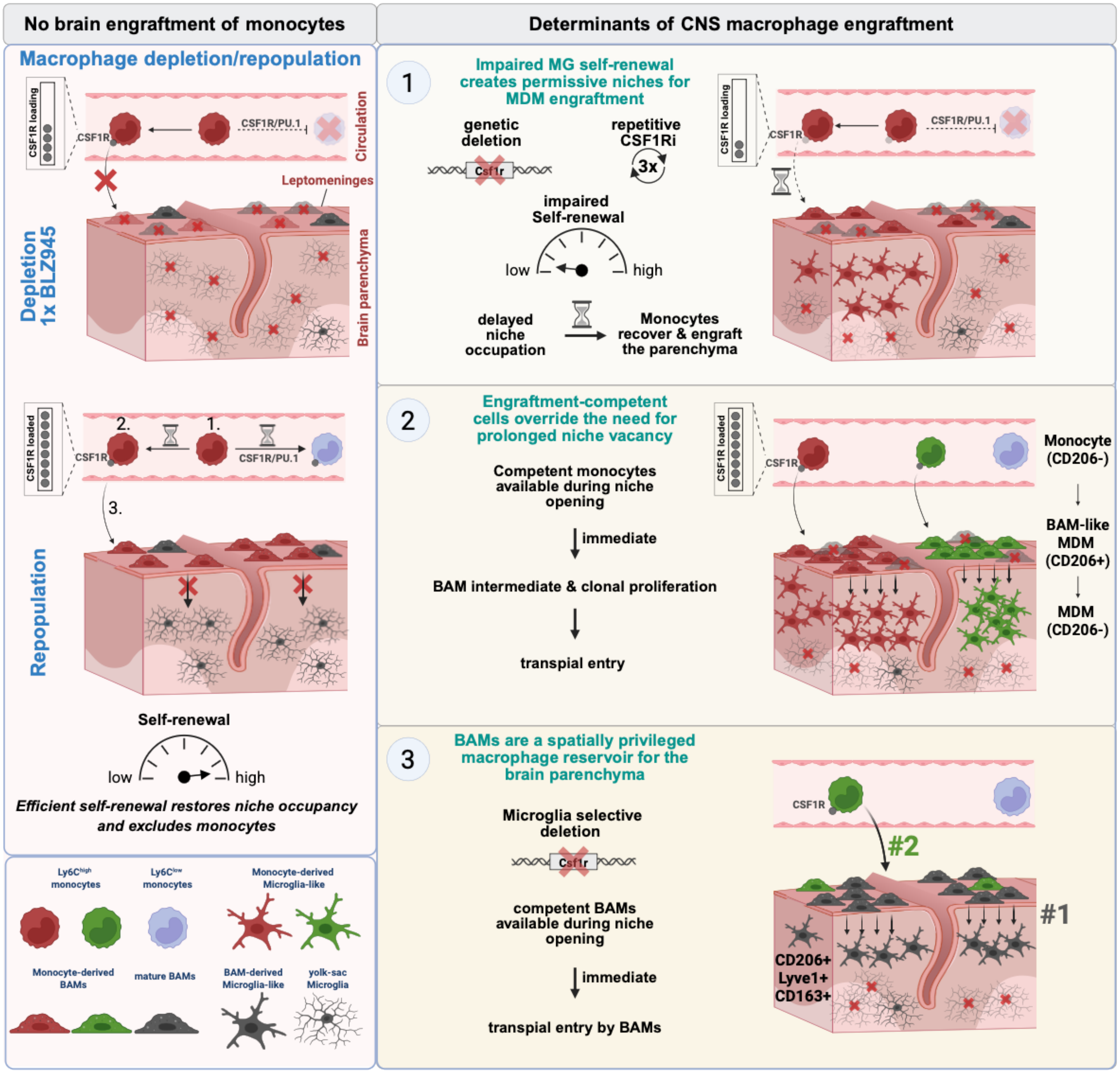



**Authors**

**Deniz Kaymak, Philip T. Williams, Sabrina Hogan, ..., Jonathan Kipnis, Rainer Glass, Gregor Hutter**

**Correspondence**

****
****



Kaymak et al. integrate complementary depletion paradigms across all CNS macrophage compartments to define the biological principles governing CNS macrophage niche maintenance and replacement. They show that tissue-resident macrophage self-renewal, CSF1R-dependent engraftment/differentiation-competence, and spatial proximity determine niche occupancy, revealing the brain-meningeal interface as a reservoir and gateway for brain macrophage maintenance and replacement in mice. Human translation of these data confirmed a border-associated macrophage (BAM)-like parenchymal macrophage signature in aged brains, independent of Alzheimer's pathology.


### Highlights

- **Exhaustion of microglia self-renewal permits peripheral macrophage engraftment into the brain.**
- **Engraftment-competent cells bypass the requirement for prolonged niche vacancy.**
- **Monocytes clonally expand in leptomeningeal and brain parenchymal niches following BAM-like transition and transpial entry.**
- **Mature BAMs directly colonize selectively vacated microglial niches**
- **Peripheral BAM-like parenchymal macrophages identified in aged human brains independent of Alzheimer's disease.**
